# The interplay between executive functions and updating predictive representations

**DOI:** 10.1101/2024.12.05.626969

**Authors:** Felipe Pedraza, Teodóra Vékony, Bence C. Farkas, Frederic Haesebaert, Romane Phelipon, Imola Mihalecz, Karolina Janacsek, Barbara Tillmann, Royce Anders, Gaën Plancher, Dezso Nemeth

## Abstract

Modifying habits, particularly unwanted behaviors, is often challenging. Cognitive research has focused on understanding the mechanisms underlying habit formation and how habits can be rewired. A key mechanism is statistical learning, the continuous, implicit extraction of probabilistic patterns from the environment, which forms the basis of predictive processing. However, the interplay between executive functions (EF) and the rewiring – or updating – of these probabilistic representations remains largely unexplored. To address this gap, we conducted an experiment consisting of four sessions: 1) Learning Phase – acquisition of probabilistic representations, 2) Rewiring Phase – updating these probabilistic representations, 3) Retrieval Phase – accessing learned representations, and 4) EF assessment, targeting five key aspects: attentional control, inhibition, working memory, flexibility, and verbal fluency. We focused on the relationship between these EF measures and the updating of previously acquired knowledge using an interindividual differences approach. Our results revealed a positive relationship between rewiring and inhibition, suggesting that better inhibitory control may facilitate the adaptive restructuring of probabilistic predictive representations. Conversely, a negative relationship was identified between rewiring and semantic fluency, implying that certain underlying aspects of verbal fluency tasks, such as access to long-term memory representations, may hinder the updating process. We interpret this relationship through the lens of competitive memory network models. Our findings indicate that the rewiring of implicit probabilistic representations is a multifaceted cognitive process requiring both the suppression of proactive interference from prior knowledge through cognitive inhibition and a strong reliance on model-free functioning.

## Introduction

Contemporary challenges like the pandemic, climate change, and rapid societal shifts result in significant environmental changes, often fostering problematic habits and maladaptive behaviors (Poldrack, 2022). In this context, the detailed comprehension of the neurocognitive dynamics underlying habit formation and the way we modify them has become a research question of the utmost importance. Traditionally, habits have been conceptualized as automatic links between stimuli and responses (Dickinson & Weiskrantz, 1997), indifferent to the outcome value of the response, which distinguishes them from goal-directed behaviors in non-human animal studies (Dickinson & Balleine, 1994). In the context of humans, habits take on a more intricate nature, defined by a set of behavioral attributes (Foerde, 2018; Seger & Spiering, 2011; Wood & Rünger, 2016). They are acquired gradually through associative learning processes over extended periods of practice, frequently without conscious awareness (Carden & Wood, 2018). Once established, these habits can be executed with minimal thought or attention, essentially operating automatically (Logan, 1988). Numerous human studies have focused on the environmental contingencies fostering successful habit updating or rewiring (Hagger et al., 2020; Verplanken et al., 2008; Verplanken & Roy, 2016; Walker et al., 2015; Wood et al., 2005). These studies commonly emphasize the modification of context cues as a primary strategy for rewiring habitual responses. Building upon the premise that habit change generally involves the strategic adjustment of environmental contingencies, we turn our attention to the neuropsychological processes that shape habit formation and modification. In this article, we focus on uncovering the internal cognitive mechanisms that govern habit formation and modification. To achieve this, we examine how interindividual differences in cognitive profiles influence habit change through the lens of a probabilistic learning task.

Statistical learning (SL) stands out as a fundamental element of the intricate process of habit formation (Graybiel, 2008; Horváth et al., 2022; Szegedi-Hallgató et al., 2017). SL refers to the cognitive function that allows the implicit extraction of probabilistic regularities from the environment, even in the absence of explicit intention, feedback, or rewards (Aslin, 2017; Conway, 2020; Kaufman et al., 2010). This cognitive mechanism plays a pivotal role in predictive processing and contributes significantly to the acquisition of diverse skills, including language (Isbilen et al., 2022; Saffran et al., 1996; Ullman, 2020), motor (Hallgató et al., 2013; Verburgh et al., 2016), musical (Rohrmeier & Rebuschat, 2012; Romano Bergstrom et al., 2012; Tillmann & McAdams, 2004), and social abilities (Baldwin et al., 2008; Parks et al., 2020; Ruffman et al., 2012). In the context of habit formation, SL fortifies the probabilistic connections between environmental cues and the invoked associated responses that constitute habitual behaviors (Hong et al., 2022; Jiang & Sisk, 2019), and facilitates the development of predictions of upcoming events, leading to shorter reaction times for expected outcomes (Tillmann & Poulin-Charronat, 2010).

For instance, consider the habitual actions of a French driver. Through SL, this driver unconsciously forms associations between the likely locations of relevant stimuli at intersections and the corresponding direction to turn their gaze. Consequently, it becomes second nature for them to instinctively look left at a crossing. Conversely, a British driver, also influenced by learned probabilistic associations, automatically looks right at a crossing (Tuhkanen et al., 2019). A potential challenge arises when a French driver, accustomed to the habitual association of relevant stimuli emerging from the left at intersections, travels to England, where relevant stimuli are more likely to appear from the right. To adapt effectively, the French driver must update or rewire the previously acquired probabilistic representations, which might prove difficult (Thompson & Sabik, 2018). Elucidating the neurocognitive processes involved in adapting to new or altered environments is crucial to understanding how probabilistic representations are updated.

Updating implicit probabilistic representations poses several challenges. Firstly, because these representations are largely unconscious, individuals may lack explicit awareness of what needs to be modified (Dresp-Langley, 2012), which can hinder rewiring. Secondly, SL has been characterized as a resilient cognitive mechanism, leading to representations that are resistant to forgetting (Kóbor et al., 2017). While this robustness is beneficial for maintaining learned associations, it can be a barrier to modifying existing representations. Thirdly, this robustness may cause proactive interference (Goedert & Willingham, 2002), where existing strong associations interfere with the encoding of novel probabilistic relationships, creating a cognitive obstacle to effectively adapt to environmental changes.

Previous research on rewiring implicit probabilistic information investigated whether providing explicit instructions could enhance the effectiveness of the rewiring process (Szegedi-Hallgató et al., 2017). While the results indicated that explicit instructions did improve rewiring, they also uncovered that the updating process could occur implicitly as well. Participants were able to adapt and rewire their implicit probabilistic representations without explicit guidance, underscoring the flexibility and adaptability of the underlying cognitive processes.

The examination of rewiring implicit probabilistic representations has also been approached through the lens of dual-process neuro-computational models of learning (Kurdi et al., 2019). These models differentiate between model-free and model-based algorithms, with the former relying solely on recent outcomes and the latter building and utilizing a model of the environment to guide choices more flexibly, but at a higher computational cost (Beierholm et al., 2011; Daw et al., 2005). The brain is postulated to alternate between these two learning systems, both within the completion of individual tasks (Doyon et al., 2009; Lee et al., 2014; Poldrack & Packard, 2003) and over the lifespan (Decker et al., 2016; Janacsek et al., 2012; Palminteri et al., 2016). In their investigation, Kurdi and colleagues employed a stimulus reevaluation protocol based on the Implicit Association Test (Greenwald & Banaji, 1995) to examine the potential for updating implicit and explicit representations via model-based and model-free learning mechanisms. Their findings showed that while the updating of explicit probabilistic representations was influenced by both model-based and model-free processes, implicit probabilistic representations were updated exclusively through model-free processes.

Cognitive control functions, also known as executive functions (EF), may be a crucial tool for effectively rewiring probabilistic representations. EF is particularly important in non-habitual situations and includes cognitive functions such as attentional control, cognitive flexibility, cognitive inhibition, and working memory updating (Diamond, 2013; Miyake et al., 2000; Miyake & Friedman, 2012). These functions are indispensable for model-based processes (Danner et al., 2011; Otto et al., 2015), and for the flexible regulation of behavior (Diamond, 2013; Gray-Burrows et al., 2019), making them pivotal in habit change (Allan et al., 2016; Allom et al., 2018). For instance, a study by Allom et al. (2018) demonstrated that cognitive remediation therapy aimed at enhancing EF could effectively disrupt bad habits associated with obesity. Investigations into the neural underpinnings of EF consistently underscore the significant involvement of the prefrontal cortex (PFC) (Badre et al., 2010; Braver et al., 2003; Funahashi & Andreau, 2013; Koechlin, 2016; Miller & Cohen, 2001; N. Osaka et al., 2004). This reliance on the PFC is further evident at an interindividual level, where superior EF is associated with greater prefrontal activity (Osaka et al., 2003), and increased functional connectivity between the PFC and other brain regions (Kondo et al., 2004). Our focus here centers on examining whether and how interindividual differences in EF may influence the rewiring of implicit probabilistic representations.

While the association between EF and habit change may suggest a positive correlation, the relationship between EF and the cognitive processes underlying habit formation is more nuanced. Greater PFC activity has been consistently shown to hinder the acquisition of implicit probabilistic information. Disruption of PFC activity through interventions such as transcranial magnetic stimulation (Ambrus et al., 2020), hypnosis (Nemeth et al., 2013), cognitive fatigue induction (Borragán et al., 2016), or dual-tasking (Filoteo et al., 2010; Smalle et al., 2022) has resulted in improved acquisition of probabilistic representations. Neuroimaging studies have revealed that decreased PFC activity (Park et al., 2022) and decreased PFC functional connectivity (Tóth et al., 2017) is also associated with enhanced SL. Similarly, at a cognitive level, studies have shown a negative relationship between EF and SL. For example, Virag et al. (2015) demonstrated that alcohol-dependent patients with weaker EF exhibited enhanced SL capacity. Similar results were observed by Pedraza et al. (2024) in a study that offered an internal replication with healthy participants. These studies suggest that while initial SL may rely on model-free learning algorithms, EF may facilitate model-based learning algorithms and align with the competition hypothesis between model-based and model-free learning. According to this hypothesis, these systems alternate in the brain during task completion, with greater reliance on one reducing reliance on the other (Daw et al., 2005; Hartley & Burgess, 2005; Poldrack & Packard, 2003). The PFC plays a crucial role in mediating between model-based and model-free systems, with the latter serving as the default but able to be overridden by cognitive control mechanisms implemented by the PFC (Lee et al., 2014; Smittenaar et al., 2013). However, it is important to note that these studies mainly focus on the relationship between cognitive control and SL during initial knowledge acquisition. The relationship between cognitive control and the updating of probabilistic knowledge remains unclear.

To our knowledge, only one study has investigated the influence of EF on the rewiring of probabilistic representations to date (Horváth et al., 2022). This study specifically examined the impact of cognitive control, as manifested by active inhibition, on rewiring. The findings revealed that participants, while actively attempting to inhibit unwanted behaviors, inadvertently impeded their ability to acquire new probabilistic knowledge. Simultaneously, they reinforced previously acquired knowledge, suggesting that cognitive control may hinder the rewiring process. However, this study did not examine how individual differences in EF capacities affect rewiring. Moreover, cognitive control functions extend beyond inhibitory control. Therefore, there remains a critical need for further investigation into how various EF interact with the rewiring of implicit probabilistic representations. Understanding these interactions is crucial for gaining insight into the neuropsychological dynamics underlying habit change.

In this study, our objective was to investigate the relationship between EF and the updating of implicit probabilistic knowledge. We aimed to characterize these cognitive processes at the interindividual level using a four-session experimental protocol comprised of empirically validated tasks and measures of EF and SL (see Methods section). In the first session, participants underwent a learning phase in which they acquired implicit probabilistic representations in a visuo-motor probabilistic sequence learning task known as the Alternating Serial Reaction Time (ASRT) task. In the second session, participants completed a rewiring phase, where structural changes to the ASRT task sequence required them to update or rewire their initially acquired probabilistic representations. On the third day, during the retrieval phase, participants were tested on both their initial and rewired knowledge. The fourth session was dedicated to assessing participants’ EF capacities using a battery of validated and reliable neuropsychological tasks targeting various aspects of cognitive control. Drawing upon the framework of competitive neurocognitive systems and existing literature, we hypothesized a negative correlation between EF capacities and the rewiring of implicit probabilistic representations.

## Methods

### Participants

Seventy-nine healthy young adults were recruited through online advertisement with eligibility based on the following criteria: participants had to be right-handed, aged under 35 years, and have no or limited musical training (less than 10 years of practice). Participants declared having no active neurological or psychiatric conditions, and not taking any psychoactive medication. Of the 79 individuals initially recruited, three did not complete the four-session experiment and were excluded from the analysis. To accurately assess the relationship between EF and rewiring performance, participants had to demonstrate acquisition of implicit probabilistic representations that could be rewired (see *Statistical analysis section for the selection criteria*). Of the 76 remaining participants, 17 did not exhibit SL during the Learning Phase. This proportion of non-learners is consistent with previous studies examining inter-individual differences in the ASRT task (Buffington et al., 2021; Farkas et al., 2024; Pedraza et al., 2024; Virag et al., 2015), addressing concerns about task sensitivity in inter-individual SL studies (Siegelman, Bogaerts, & Frost, 2017; Siegelman, Bogaerts, Christiansen, et al., 2017). Thus, the data from the remaining 59 participants (32 females, M _age_ = 22.51 years; SD _age_ = 2.92 years; M _education_ = 15.34 years; SD _education_ = 1.64 years) were included in the analyses. Further details on participant characteristics can be found in Supplementary Table 1. All participants provided signed informed consent agreements and received a 200 euro financial compensation for their participation in the four sessions of the experiment. The relevant institutional review board (i.e., the “Comité de Protection des Personnes, CPP Est I” ID: RCB 2019-A02510-57) gave ethical approval for the study.

### Tasks

#### Measure of SL and rewiring: The Alternating Serial Reaction Time (ASRT) task

To evaluate the acquisition and rewiring of implicit probabilistic information, we utilized a visuomotor SL task that has been empirically validated (Buffington et al., 2021) and previously established as reliable (Farkas et al., 2024), known as the Alternating Serial Reaction Time (ASRT) task (Howard & Howard, 1997). Employing a modified version of the ASRT task (Figure 1A), our study adhered to a structured three-session paradigm (Szegedi-Hallgató et al., 2017), designed to investigate the phases of acquisition, rewiring, and consolidation of implicit probabilistic information.

**Figure 1.**
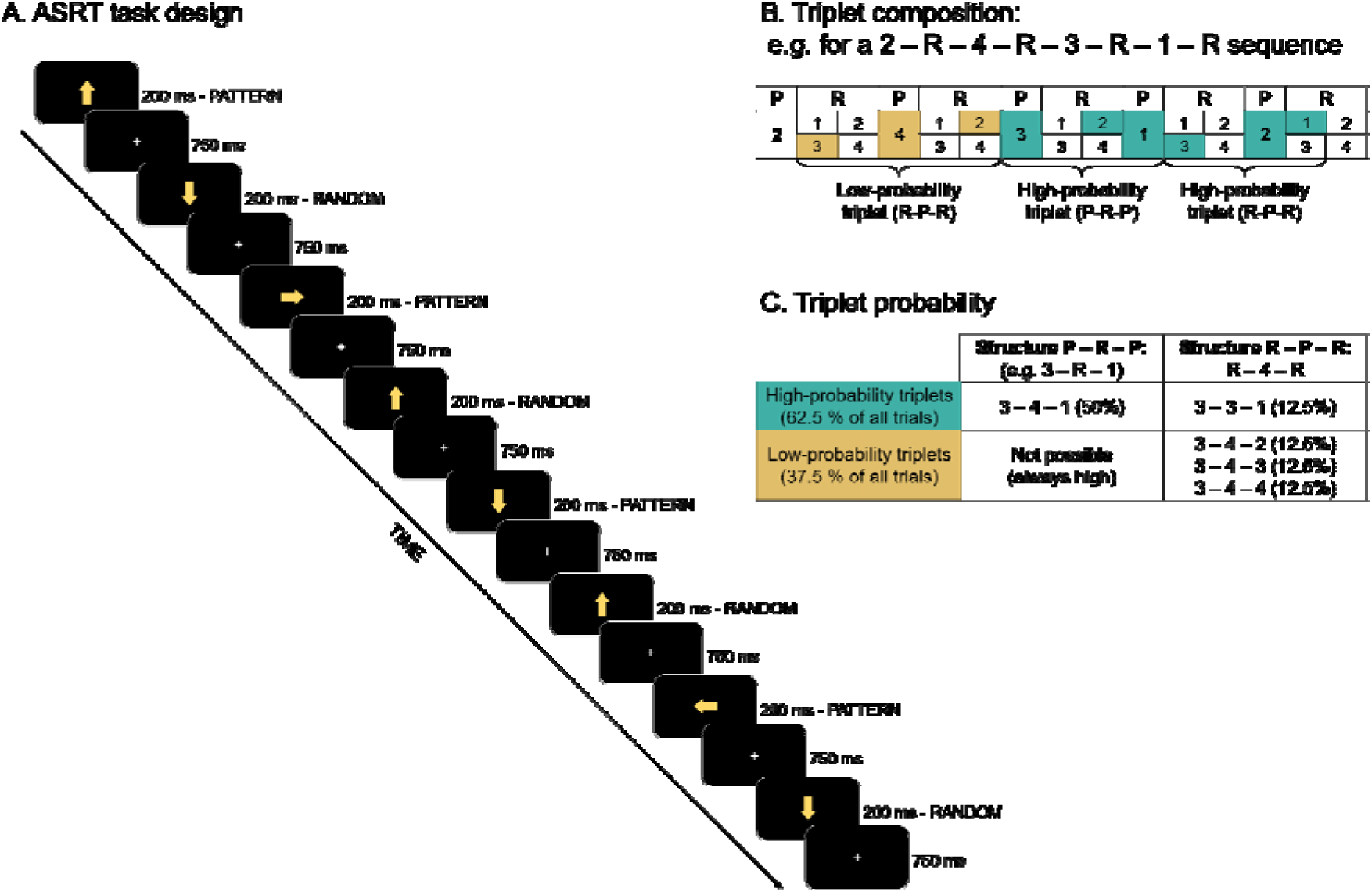
ASRT task structure. A) ASRT task design. The presentation stimuli (yellow arrows, indicating one of the four cardinal directions), followed an eight-element sequence alternating between pattern (P) and random (R) stimuli. This sequence was presented 10 times per block. Here we illustrate an example of a 2-R-4-R-3-R-1-R sequence. **B) Triplet composition**. Each stimuli position can be coded with a number. Here we have 1=left, 2=up, 3=down, 4=right. From the alternating structure of the sequence between P and R stimuli, certain triplets (sequences of three consecutive stimuli) have a higher probability of occurrence than others. For example, in the 2-R-4-R-3-R-1-R sequence, the triplet 3-2-1 has a higher probability of occurrence since it can be found in P-R-P and R-P-R structures. The triplet 3-4-2 has a lower probability of occurrence since it can only appear in R-P-R structures. igh-probability triplets could end in either a pattern or random element, while low-probability triplets always concluded with a random element. High- and low-probability triplets are denoted in green and yellow, respectively. **C) Triplet probability.** High-probability triplets account for 62.5% of all trials and low-probability triplets account for the rest. Notably, for each high-probability triplet (e.g., 3-4-1), there are low-probability triplets with three different last elements (e.g., 3-4-2, 3-4-3 and 3-4-4). This structure overall results in each high-probability triplet being five times more likely to occur than the low-probability triplets.

#### Session 1: Learning phase

During the Learning Phase, participants performed Sequence A of the ASRT task. In each trial, a yellow arrow appeared at the center of the screen for 200 ms, pointing in one of four possible directions (left, up, down, or right). This was followed by a fixation cross displayed for 500 ms. Participants were instructed to respond as quickly as possible by pressing the button corresponding to the arrow’s direction on a Cedrus RB-530 response box. Finger placement was controlled as follows: the left index finger for the up button, the right thumb for the down button, the right index finger for the right button, and the left thumb for the left button. Correct responses were followed by a 750 ms display of the fixation cross, while no response or incorrect responses were met with a 500 ms display of an exclamation mark or an “X,” respectively, followed by a 250 ms fixation cross. Each block consisted of 85 stimuli, with the initial five presented randomly for practice, followed by 10 repetitions of the eight-element alternating sequence. The Learning Phase comprised 25 blocks of the ASRT task.

Unbeknownst to participants, the stimuli followed a structured sequence alternating between predefined pattern elements (P) and random elements (R) (e.g., 2-R-4-R-3-R-1-R, where the numbers represent the direction of the yellow arrows: 1 for left, 2 for up, 3 for right, 4 for down, and “R” indicating a randomly chosen direction out of the four possibilities). This structure resulted in certain triplets (runs of three trials) having a higher probability of occurrence than others (Figure 1B). For instance, in the 2-R-4-R-3-R-1-R sequence, triplets like (2-3-4) or (3-4-1) were more likely to occur than triplets like (2-3-2) or (4-3-1) since the former triplets could appear in both P-R-P or in R-P-R structures whereas the latter triplets could only appear in R-P-R structures (Figure 1C). Consequently, the former triplets were five times more likely to occur than the latter, hence they were categorized as high-probability (H) and low-probability (L) triplets, respectively. Previous studies using the ASRT task consistently demonstrate that, with practice, participants respond more quickly to the last element of high-probability triplets compared to low-probability triplets, indicating SL (Janacsek et al., 2012; Janacsek & Nemeth, 2012; Kóbor et al., 2017). Importantly, participants typically do not develop explicit knowledge of the sequence structure despite this learning effect (Janacsek & Nemeth, 2012; Kóbor et al., 2017; Vékony et al., 2022). Thus, learning in the task is assessed by calculating the reaction time difference between the last elements of high-probability and low-probability triplets, and this learning measure reflects implicit SL.

#### Session 2: Rewiring Phase

During the Rewiring Phase, participants completed 25 blocks of Sequence B of the ASRT task, which involved structural modifications to the original sequence of the Learning Phase (Figure 2A). Specifically, the last two pattern elements (P) of the original eight-element sequence were inverted. For example, if Sequence A was structured as 2-R-4-R-3-R-1-R, Sequence B became 2-R-4-R-1-R-3-R. This change in sequence structure affected the occurrence probability of certain triplets (Figure 2B). Specifically, 75% of triplets classified as high-probability during the Learning Phase became low-probability in the Rewiring Phase (HL, with the first letter denoting the triplet probability in Sequence A and the second letter referring to the probability in Sequence B). Conversely, new high-probability triplets (LH) emerged during the Rewiring Phase, replacing those that became low-probability. Meanwhile, the probability of other triplets remained constant, either maintaining a low probability (LL) or a high probability (HH) in both phases.

**Figure 2.**
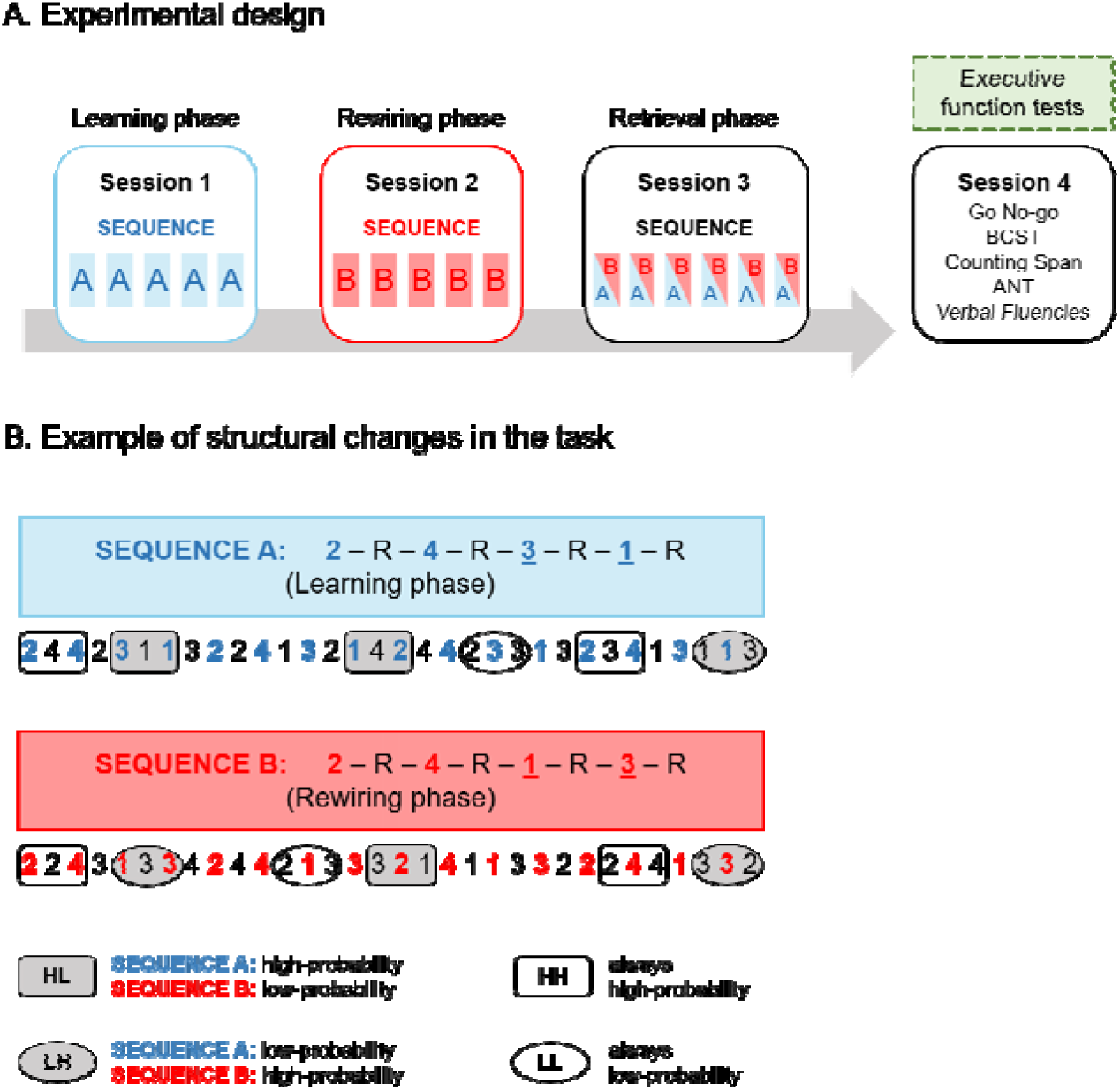
Experimental design and sequence changes. A) Experimental design. The experiment comprised four sessions: the first three were held 24 hours apart, and the fourth was independent. Sessions one to three included Learning, Rewiring, and Retrieval phases. In the Learning Phase, participants completed 25 blocks of Sequence A of the Alternating Serial Reaction Time (ASRT) task to acquire initial probabilistic representations. During the Rewiring Phase, structural changes introduced Sequence B, and participants updated their representations over 25 ASRT blocks. The Retrieval Phase involved alternating between Sequences A and B every five blocks for 30 blocks, probing knowledge of both sequences. The starting order of the sequences in the Retrieval Phase was counterbalanced across participants (half started with Sequence A and the other half started with Sequence B). In the figure, blue, red, and half-colored rectangles represent a bin (five blocks) of the ASRT task. The fourth session included executive functions (EF) assessments with five tasks: GoNoGo, Berg Card Sorting Test (BCST), Counting Span, Attention Network Test (ANT), and three Verbal Fluency tasks (action, semantic, and lexical fluency). **B) Example of structural changes in the ASRT task.** The figure shows the stimulus order for Sequences A and B, coded as numbers, with predetermined stimuli in color alternating with random stimuli (R in black). The alternating structure resulted in high- and low-probability triplets. The Rewiring Phase modified Sequence A into Sequence B, altering triplet probabilities. Specifically, 75% of initially high-probability triplets became low-probability (HL trials, gray squares), replaced by new high-probability triplets that were initially low-probability in Sequence A (LH trials, gray ovals). Other triplets maintained their original probability, being either low-probability (LL trials, white ovals) or high-probability (HH trials, white squares) in both phases.

The LH triplets were crucial for assessing the acquisition of new knowledge during the Rewiring Phase. As these triplets transitioned from low- to high-probability in the Rewiring Phase, any associated knowledge could be considered acquired during this session. The LL triplets, on the other hand, served as a baseline for controlling general practice effects. Thus, the reaction time difference between LH and LL triplets was used as a key measure of participants’ rewiring ability in the second session (Horváth et al., 2022).

#### Session 3: Retrieval Phase

During the Retrieval Phase, participants completed 30 blocks of the ASRT task. Unbeknownst to them, they were tested on both Sequence A and Sequence B. To ensure balanced testing, half of the participants started with Sequence A followed by Sequence B, while the other half started with Sequence B followed by Sequence A. Participants completed five blocks of one sequence before switching to the other sequence, and this alternation continued until all participants completed 15 blocks of each sequence (i.e., 30 blocks in total). Knowledge of Sequence A was assessed by computing the difference between HL and LL triplets, whereas knowledge of Sequence B was evaluated by comparing LH and LL triplets (Horváth et al., 2022).

#### Measures of executive functions (Session 4)

The assessment of EF was guided by the Unity and Diversity model of EF, which encompasses inhibition, updating, and shifting processes (Miyake et al., 2000; Miyake & Friedman, 2012). Additionally, we examined executive control of attention, working memory, and verbal fluency, which are also prefrontal-dependent functions (Curtis & D’Esposito, 2003; N. Osaka et al., 2004; Paneri & Gregoriou, 2017; Phelps et al., 1997; Robinson et al., 2012; Rossi et al., 2009). To operationalize these constructs, we employed a battery of well-established and reliable cognitive tests, which included the Attentional Network Test, the Berg Card Sorting Test, the Counting Span Task, the GoNoGo Task, and three verbal fluency tasks. These tests were selected based on their well-documented validity and reliability in previous research (Conway et al., 2005; Fox et al., 2013; Harrison et al., 2000; Langner et al., 2023; Piper et al., 2015; Woods et al., 2005), ensuring robust measurement of the targeted cognitive functions.

#### Attentional network test (ANT)

The ANT assessed three distinct attention networks: the alerting, orienting, and executive networks (Fan et al., 2002). Participants had to identify whether a central arrow, embedded in an array of five arrows, pointed left or right. The array could appear above or below a fixation cross and might be preceded by a spatial cue indicating the subsequent location. Additionally, in half of trials, the central arrow was congruent with the other arrows (pointing in the same direction), and in the other half it could be incongruent with the other arrows (pointing in the opposite direction).

To compute network scores, we adhered to the standard methodology (Fan et al., 2002). The alerting component was derived by subtracting the mean reaction time (RT) of the central cue conditions from the mean RT of the no-cue conditions for each participant. The orienting component was computed by subtracting the mean RT of the spatial cue conditions from the mean RT of the central cue conditions. The executive component of attention was determined by subtracting the mean RT of all congruent conditions from the mean RT of all incongruent conditions. In this task, higher scores across the alerting, orienting, and executive domains indicated enhanced performance in the respective aspects of attention. In simpler terms, elevated scores suggest better attentional abilities in these three facets of attentional processing.

#### Berg card sorting task (BCST)

We assessed set shifting or cognitive flexibility using the computerized version of the Berg Card Sorting Test (Berg, 1948) with 64 cards (BCST.64), available in the Psychology Experiment Building Language (PEBL) software (Mueller & Piper, 2014a). In this test, a set of four cards with varying characteristics—color, shape, and number of items—was displayed at the top of the screen. Participants were instructed to match new cards to those at the top based on one of the three characteristics, without knowing the current rule but received feedback after each attempt. The rule could change during the task. Cognitive flexibility was measured through perseverative errors, which indicated failure to adapt quickly after a rule change. In essence, these errors provide insights into the participant’s ability to adjust their cognitive approach when the task requirements change.

#### Counting Span (CSPAN) task

To assess updating or working memory capacity, we used the CSPAN task (Case et al., 1982). In this task, various shapes, including blue circles, blue squares, and yellow circles, were presented on a computer screen. Participants were instructed to audibly count and remember the number of blue circles (targets) among other shapes (distractors) in a series of images. Each image consisted of three to nine blue circles, one to nine blue squares, and one to five yellow circles. After each trial, participants verbally reported the total number of targets in the image. If the count was accurate, the experimenter proceeded to the next trial. A recall cue at the end of a set prompted participants to recall the number of targets for each image in their order of presentation. If the recall was correct, a new set with an additional image commenced, up to maximum six images in a set. If a participant made an error in recall, the task was halted, and a new run, starting with a set of two images, commenced. Each participant completed three runs of the task. Memory span was computed as the mean of the highest set sizes that the participant correctly recalled in the three runs. This measure provides an insight into the participant’s ability to efficiently update and maintain information in working memory.

#### GoNoGo (GNG) task

We measured cognitive inhibition with the computerized version of the GNG task available in the PEBL software (Mueller & Piper, 2014b). In the GNG task, participants were instructed to respond to certain stimuli (“go” stimuli) by clicking on a button as fast as possible and to refrain from clicking on other stimuli (“no-go” stimuli). In this version, participants were presented with a 2×2 array with four blue stars (one in the center of each square of the array). Every 1500 ms, a stimulus (the letter P or R) would appear for 500 ms in the place of one of the blue stars. For the first half of the task, the letter P would be the “go” stimulus and the letter R would be the “no-go”. This rule would be then inverted in the second half of the task. Participants completed 320 trials. The ratio between “go” and “no-go” trials was 80:20, respectively. Cognitive inhibition capacity in the GNG task was measured by the d’:

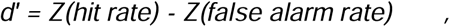

with higher *d’* scores indicating better cognitive inhibition.

#### Verbal fluency tasks

Verbal fluency was assessed through three subtasks, examining the lexical, semantic, and action components of verbal fluency. In the lexical fluency subtask, participants were tasked with generating as many words as possible beginning with the letter “P,” a common practice in the French version of phonemic fluency (St-Hilaire et al., 2016). For the semantic fluency subtask, participants were instructed to name animals, and for the action fluency subtask, they had to articulate isolated verbs describing actions feasible for a person to perform.

In each subtask, participants aimed to produce as many words as possible within one minute while adhering to specific guidelines: avoiding word repetitions, words with the same etymological root, and proper nouns. Any violation of these rules constituted an error. The score for each verbal fluency subtask was calculated by subtracting the number of errors from the total words generated within the one-minute timeframe. A higher score in each verbal fluency component denoted greater verbal fluency capacity, reflecting the participant’s ability to generate words efficiently while adhering to specified constraints.

### Experimental Design

Participants took part in a four-session experiment (Figure 2A). The first three sessions were dedicated to assessing the acquisition, rewiring, and retrieval of implicit probabilistic information, each separated by a 24-hour interval. In the initial session (Learning Phase), participants’ statistical learning (SL) abilities were evaluated using the ASRT task. Here, participants acquired associations related to Sequence A. In the second session (Rewiring Phase), a structural change was introduced to the task by incorporating Sequence B (see Tasks section for details). This modification allowed us to measure participants’ rewiring abilities. In the third session (Retrieval Phase), participants were tested on both sequences A and B in a counterbalanced order, providing insight into the consolidation of initial and rewired knowledge. Half of the participants started the Retrieval Phase with Sequence A and the other half with Sequence B. The fourth session was conducted independently from the first three sessions and was planned according to each participant’s availability. In this session, prefrontal-dependent cognitive functions were assessed with a battery of well-established and reliable neuropsychological tasks: the ANT, the BCST, the CSPAN task, the GNG task and three variants of the Verbal Fluency tasks (action, lexical and semantic fluency).

### Statistical analysis

Evaluating someone’s ability to rewire their knowledge during the Rewiring Phase requires that the initial knowledge is acquired during the Learning Phase. Therefore, participants who did not exhibit initial SL during the Learning Phase (i.e., L-H<0) were excluded. This refinement in participant selection aimed to enhance the relevance and meaningfulness of all subsequent analyses focusing on the relationship between rewiring and EF. Seventeen participants did not demonstrate SL during the Learning Phase. Consequently, our analyses were performed using the data of the remaining 59 participants.

#### Learning, rewiring and retrieval trajectories

For the Learning, Rewiring, and Retrieval Phases, the first five trials (comprising five warm-up random trials) of each block were excluded from the analysis, along with all incorrect responses throughout all trials. Additionally, trials featuring trills (e.g., 2-1-2) or repetitions (e.g., 2-2-2) were omitted from the analysis because participants tend to respond more rapidly to these types of triplets due to pre-existing tendencies (Song et al., 2007). To facilitate the analysis and improve the signal to noise ratio, the blocks of the ASRT task were organized into units of five blocks, referred to as bins.

In the first session (Learning Phase), learning performance on Sequence A was assessed by calculating the median reaction times (RTs) within each bin for correct responses in high-probability triplets specific to Sequence A (HL) and low-probability triplets common to all sessions (LL), which served as a baseline to control for general practice effects. Subsequently, learning scores were computed for each bin by subtracting the median RTs of HL triplets from the median RTs of LL triplets. A higher difference score between HL and LL triplets indicated better initial learning of probabilities. The learning scores for each bin were then averaged to generate a single score reflecting statistical learning (SL) for each participant in the Learning Phase.

In the second session (Rewiring Phase), rewiring performance on Sequence B was evaluated by calculating the median RTs for correct responses in high-probability triplets specific to Sequence B (LH) and LL triplets. Data processing followed a similar approach to the first session, with the unique distinction being that rewiring scores resulted from the difference between LH and LL triplets. A higher difference score between LH and LL triplets indicated better rewiring performance.

During the third session (Retrieval Phase), participants alternated between bins of sequences A and B in a counterbalanced order. Data for assessing the retrieval of Sequence A was processed similarly to the Learning Phase, while data for Sequence B retrieval followed the same procedure as in the Rewiring Phase.

To evaluate SL in the Learning and Rewiring Phases separately, we conducted a repeated measures analyses of variance (RM ANOVAs) with two factors for each phase: bin (1-5) and triplet (HL vs. LL for the Learning Phase; LH vs. LL for the Rewiring Phase). To compare the retrieval of Sequences A and B in the third session, we conducted an RM ANOVA with two factors: bin (1-3; corresponding to the three bins presented for each sequence in the Retriaval Phase), and sequence (A vs. B). In all ANOVAs, Greenhouse-Geisser epsilon correction was used when Mauchly’s test of sphericity indicated that sphericity could not be assumed. Original df values and p values are reported together with partial eta-squared (η_p_^2^) as the measure of effect size.

To complement these analyses, we conducted Bayesian RM ANOVAs using JASP (JASP Team, 2019). Bayesian analysis allows us to provide evidence in favor of the null hypothesis, rather than failing to reject it. The inclusion Bayes factor quantifies the change from prior inclusion odds to posterior inclusion odds and can be interpreted as evidence in the data for including a predictor or factor. More generally, Bayes factors quantify the relative weight of evidence provided by the data for two theories, the null and the alternative hypotheses, H0 and H1 (Dienes, 2014). According to a commonly used classification (Wagenmakers et al., 2011), BF values between 1 and 3 indicate anecdotal evidence, values between 3 and 10 indicate substantial evidence, and values above 10 indicate strong evidence for H1. Conversely, values between 1/3 and 1 indicate anecdotal evidence, values between 1/10 and 1/3 indicate substantial evidence, and values below 1/10 indicate strong evidence for H0. Values near 1 indicate that the data do not favor either hypothesis.

#### Correlations

For our correlation and multiple regression analyses, we derived single indices of learning, rewiring, and retrieval of sequences A and B by averaging scores across all bins for each participant. Prior research has indicated that while the reliability of individual bins is typically low, averaging across at least five bins results in acceptable levels of reliability (Farkas et al., 2024). Given the importance of using reliable measures for assessing inter-individual differences, our analysis of the relationship between rewiring and EF utilized these average scores. Our reported Pearson’s correlations between learning, rewiring, retrieval and EF scores include Benjamin-Hochberg adjusted p-values to account for multiple comparisons. Additionally, we calculated Bayes factors using JASP (JASP Team, 2019) with default priors (stretched beta distribution with a width of 1) to further validate our findings.

#### Multiple regression analysis

To assess the relationship between EF and the different phases of the experiment (i.e., learning, rewiring, and retrieval), we conducted multiple linear regression analyses. The predictor variables included the scores of the executive component of the ANT, BCST, CSPAN task, GNG task, and verbal fluency tasks. Note that the alerting and orienting components of the ANT were not included in the models as these do not reflect EF (Fan et al., 2002, 2005). The regression analyses were applied systematically to the outcome variables for each experimental phase.

For the Learning Phase, the EF tests were used as predictor variables to assess their contribution to the variance in the initial learning performance on (Sequence A). For the Rewiring Phase, we explored the association between EF test scores and rewiring performance (on Sequence B). For the Retrieval Phase, two separate multiple regression analyses were conducted to assess the relationship between EF and the retrieval of Sequence A and Sequence B. Multicollinearity among predictor variables was assessed using variance inflation factor (VIF) in each model. All VIF values were below the commonly accepted threshold of 5, with the highest value being 1.804 for action fluency, indicating that multicollinearity is not a concern in our models.

## Results

### Learning, rewiring and retrieval trajectories

#### Learning phase

To evaluate SL in the learning phase, we conducted a repeated measures analysis of variance (RM ANOVA) with two factors: bin (1-5) and triplet (HL vs. LL). The main effect of bin was significant (*F*_(4, 232)_ = 13.056, *p* < .001, η_p_^2^ = .184, BF_inclusion_L=3.527e+9), indicating general skill learning, i.e., participants became faster overall as the task progressed, irrespective of triplets (bin 1 significantly slower than all further bins all *p*-values < .001, all other pairwise comparisons *p* > .248). The main effect of triplet was significant (*F*_(1, 58)_ = 12.395, *p* < .001, η_p_^2^ = .176, BF_inclusion_L= 14816.224), with average responses to HL triplets being faster than responses to LL triplets (M_HL_ = 358 ms, 95% CI = [352, 364]; M_LL_ = 362 ms, 95% CI = [356, 368]; d = .157), revealing successful SL during the Learning Phase (Figure 3A). The bin by triplet interaction was non-significant (*F*_(4, 232)_ = 1.644, *p* = .164, η_p_ ^2^ = .028, BF_inclusion_L= .121) indicating that SL did not change substantially across the bins.

**Figure 3.**
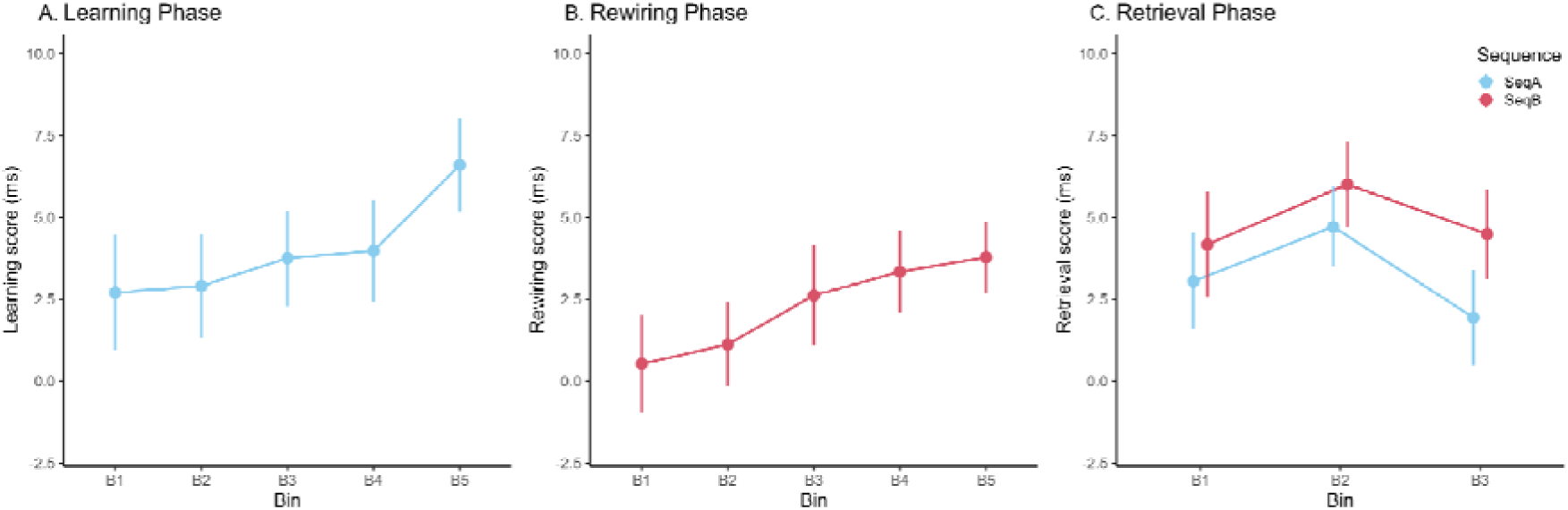
Acquisition and retrieval of implicit knowledge during the different phases of the experiment. **A) Leaning Phase.** Acquisition of the initial sequence during the Learning Phase. The learning scores for each bin denote the difference of RTs between LL and HL triplets. Participants acquired the probabilistic regularities of Sequence A, responding faster to HL triplets compared to LL triplets. **B) Rewiring Phase.** Acquisition of a new sequence during the Rewiring Phase. The rewiring scores for each bin denote the difference of RTs between LL and LH triplets. Participants acquired the probabilistic regularities of Sequence B, responding faster to LH triplets compared to LL triplets. **C) Retrieval Phase.** Retrieval of both sequences during the Retrieval Phase. The retrieval scores denote the differences between HL and LL triplets for Sequence A (in blue), and LH and LL triplets for Sequence B (in red). Sequence A and Sequence B knowledge was retrieved equivalently and did not vary across the bins. Error bars in all figures denote the standard error of the mean.

#### Rewiring phase

To evaluate SL in the Rewiring Phase, we conducted an RM ANOVA with two factors: bin (1-5) and triplet (LH vs. LL). The main effect of bin was non-significant (*F*_(4, 232)_ = 0.513, *p* = .726, η_p_^2^ = .009, BF_inclusion_ =.006), indicating that, irrespective of triplets, the RTs did not progress in the second session. Importantly, the main effect of triplet was significant (*F*_(1, 58)_ = 5.596, *p* = .021, η_p_ ^2^ = .088, BF_inclusion_ = 12.602), with average responses to LH triplets being faster than responses to LL triplets (M_LH_ = 347 ms, 95% CI = [341, 353]; M_LL_ = 349 ms, 95% CI = [344, 355]; d = .098), indicating updating of the initial knowledge during the second session (Figure 3B). The bin by triplet interaction was non-significant (*F*_(4, 232)_ = 1.797, *p* = .130, η_p_^2^ = .030, BF_inclusion_L= .001), indicating that rewiring performance did not vary substantially across the bins.

#### Retrieval phase

To evaluate the retrieval of sequences A and B in the third session, we conducted an RM ANOVA, this time on the learning scores, with two factors: bin (1-3) and sequence (A vs. B). Learning scores for both sequences were significantly different from 0 in almost all bins (*p*s < .043), except the Sequence A learning score for bin 3, which did not significantly differ from 0 (*p* = .190). The main effect of bin was non-significant (*F*_(2, 116)_ = 2.081, *p* = .129, η_p_ ^2^ = .035, BF_inclusion_C:=.168) indicating that learning scores remained relatively constant during the retrieval phase. The main effect of sequence was non-significant (*F*_(1, 58)_ = 3.220, *p* = .078, η_p_^2^ = .053, BF_inclusion_C:=.421) indicating that both sequences were retrieved equally during the third session (Figure 3C), although there is a trend towards smaller Sequence A learning scores (M_SeqA_ = 3.23 ms, 95% CI = [1.02, 5.45]; M_SeqB_ = 4.89 ms, 95% CI = [2.63, 7.15]; d = .152). The bin by sequence interaction was non-significant (*F*_(2, 116)_ = 0.300, *p* = .742, η_p_ ^2^ = .005, BF_inclusion_C:= .022) indicating that the retrieval of sequences A and B did not vary across the bins.

### Correlations

Bivariate Pearson’s correlations with Benjamin-Hochberg adjusted p-values between all variables are presented in Figure 4. Overall, most EF measures showed weak or non-significant correlations with learning, rewiring, and retrieval scores for sequences A and B. The strongest correlations were found between the rewiring and GNG scores (Pearson’s r = .439, 95% CI = [.206, .625], *p* = .003, BF_10_C:= 59.484) and between the rewiring and Semantic fluency scores (Pearson’s r = −.378, 95% CI = [−.578, −.135], *p* = .0151, BF_10_C:= 11.450).

**Figure 4.**
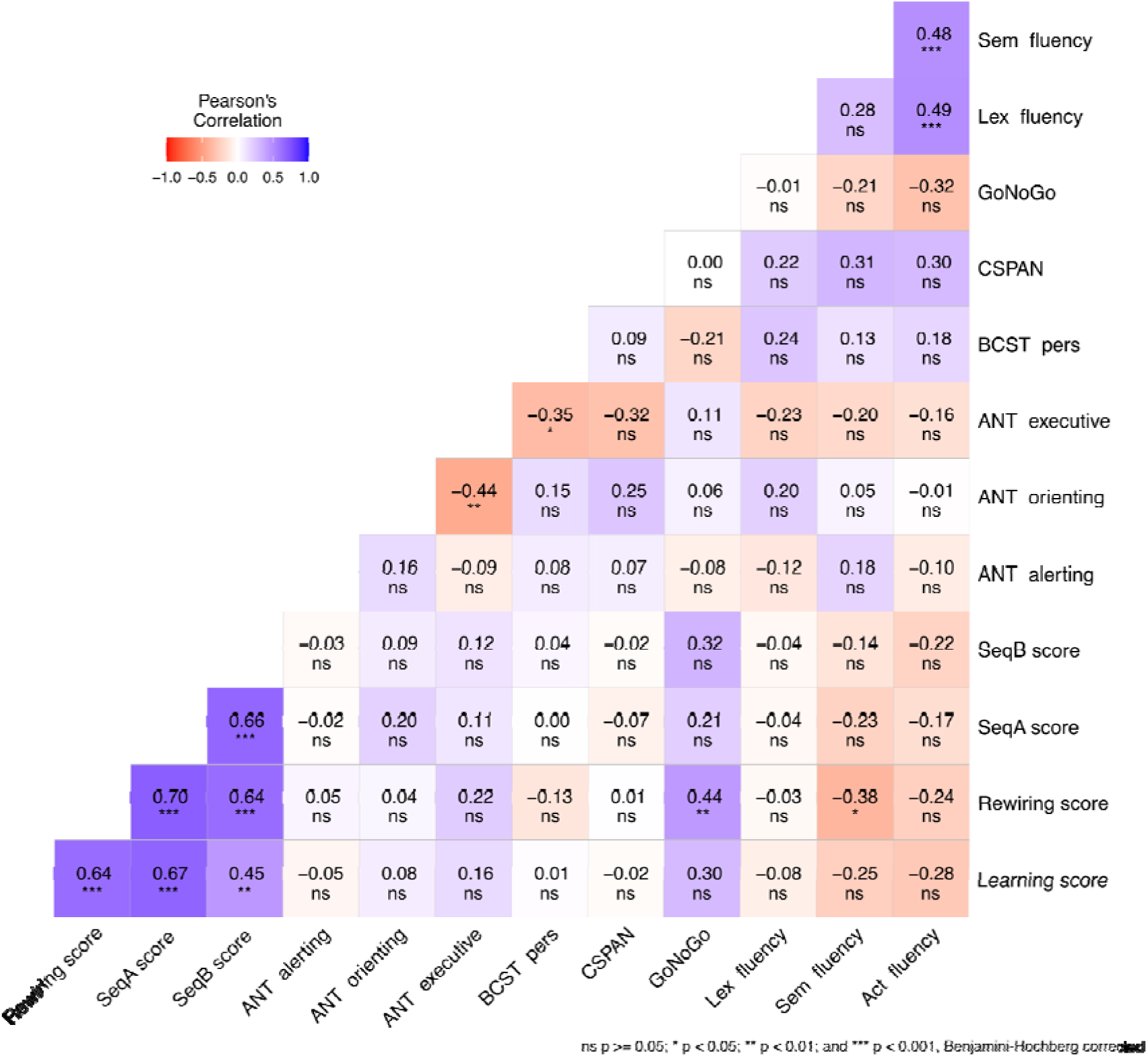
Correlations Between Learning, Rewiring, Retrieval Scores, and EF Measures. Negative and positive correlations are visually represented by red and blue backgrounds, respectively. A significant positive correlation was found between cognitive inhibition (measured by GNG) and rewiring performance, while a significant negative correlation was found between semantic fluency and rewiring. Benjamin-Hochberg corrected p-values are included in the matrix. The Learning score reflects knowledge acquired in Session 1, while the Rewiring score reflects the updating of knowledge (i.e., rewiring performance) in Session 2. SeqA score indicates the retrieval of initial knowledge in Session 3, and SeqB score reflects the retrieval of updated knowledge in Session 3. ANT alerting, orienting, and executive represent the three aspects of the Attentional Network Test. BCST pers score denotes cognitive flexibility performance in the Berg Card Sorting Test. CSPAN indicates working memory performance in the Counting Span task. GoNoGo reflects inhibition performance in the GoNoGo task. Lastly, Act, Sem, and Lex fluency scores represent performances in the Action Fluency, Semantic Fluency, and Lexical Fluency tasks, respectively.

### Multiple regression analysis

#### Rewiring phase

To test whether EF scores predict the rewiring of implicit probabilistic information, a multiple regression analysis was conducted with the rewiring score as the dependent variable. Predictor variables included the executive component of ANT, BCST, CSPAN, GNG, and verbal fluency tasks (action, semantic, and lexical). Together, these predictors explained 22.6% of the variance in rewiring performance (Adjusted R^2^ = .226, *F*_(7, 51)_ = 3.418, *p* = .005). Notably, the GNG score emerged as a strong predictor of rewiring (B = 5.881, 95% CI = [1.458, 10.304], *p* = .010, β = .341), suggesting that better inhibition, as measured by GNG performance, was associated with more effective rewiring of implicit probabilistic representations (Figure 5A). In contrast, the semantic fluency score negatively predicted rewiring performance (B = −0.455, 95% CI = [−0.838, −0.073], *p* = .021, β = −.325), indicating that better semantic fluency performance was inversely related to rewiring (Figure 5B). Other EF scores, including ANT executive, BCST, CSPAN, lexical fluency, and action fluency, did not significantly contribute to the model (Supplementary Table 2).

**Figure 5.**
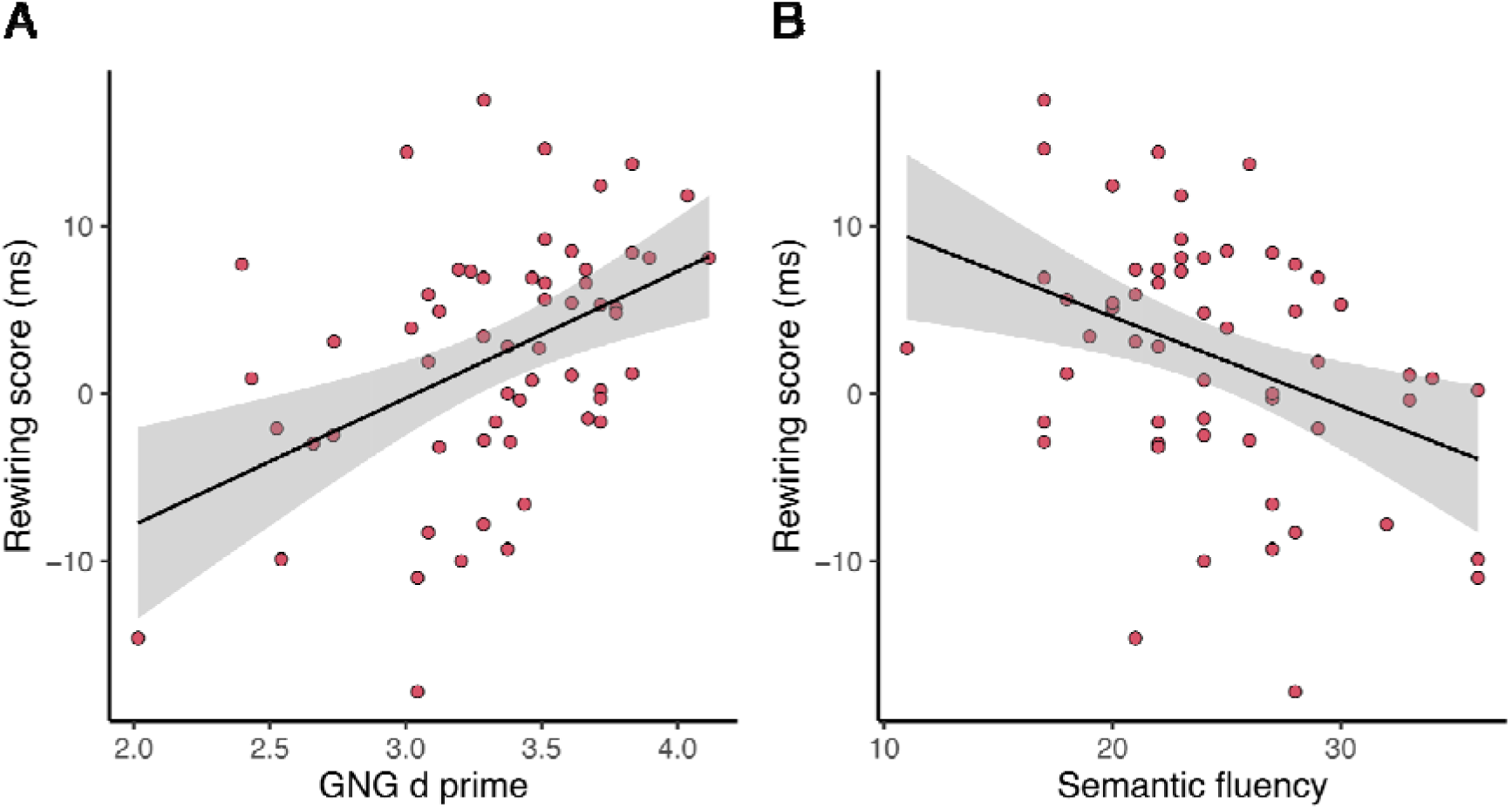
Relationship between the Rewiring score and the GNG d-prime and Semantic fluency scores. Each data point represents a participant. The black line represents the best-fit regression line, and the gray shading indicates the 95% confidence interval.

#### Learning and Retrieval phases

In line with the correlation results reported above, multiple regression analyses for learning in Session 1 and retrieval of Sequences A and B in Session 3 revealed no significant predictors. Details of these multiple regression results are presented in Supplementary Tables 3, 4, and 5, respectively.

## Discussion

When the environment changes, our old habits and predictions no longer function optimally. Our study aimed to explore how individual differences in EF affect the updating of our predictive models, leading to changes in our habits. Our findings revealed distinct correlations between EF and the rewiring of implicit probabilistic representations. Specifically, we observed a positive relationship between inhibition, measured by the GoNoGo task, and rewiring. We also observed a negative relationship between semantic fluency and rewiring. These correlations were further confirmed by multiple regression analysis, which showed that while GoNoGo scores positively predicted rewiring performance, semantic fluency scores negatively predicted it. This suggests that individuals with stronger inhibitory control and/or lower semantic fluency tend to exhibit a better capacity to update their probabilistic representations during rewiring.

The positive relationship between rewiring and the GoNoGo task suggests that the ability to inhibit irrelevant or outdated information and behaviors may facilitate the adaptation to new probabilistic contingencies in the presence of pre-existing knowledge. This positive correlation resonates with existing literature on EF and habit change, where inhibitory control has been shown to play a crucial role in disrupting and modifying unhealthy habits and ingrained behaviors (Allom et al., 2016, 2018; Costa et al., 2019; Galla & Duckworth, 2015; Preuss et al., 2017). Cognitive inhibition may facilitate habit change by shielding the individual from maladaptive memory representations (Danner et al., 2011) and suppressing automatic motor responses tied to old habits (Jahanshahi et al., 2015).

Intriguingly, this finding contrasts with previous research, such as Horvath et al. (2022), which reported that inhibitory control was detrimental to the rewiring of probabilistic representations. Notably, efforts to understand the relationship between SL and EF have led to various experimental approaches, generally categorized into four main types. The first involves behavioral studies adopting an interindividual differences approach, measuring SL and EF with separate tasks and investigating the correlations between their performance indices. These studies have found both positive and negative associations, suggesting potential cooperation, competition, and instances of independence (Park et al., 2020; Pedraza et al., 2024; Virag et al., 2015). The second category includes interventional studies that deplete, disrupt, or suppress EF to observe how this affects SL, generally finding a competitive relationship, with weakened EF leading to better SL performance (Borragán et al., 2016; Nemeth et al., 2013; Smalle et al., 2022). The third category focuses on brain network studies, examining the relationship between SL and the disengagement of brain networks supporting EF, and finding that such disengagement improves SL (Ambrus et al., 2020; Park et al., 2022; Tóth et al., 2018). The fourth category employs combined paradigms that integrate features of both SL and EF in a single experimental design, providing direct insight into their real-time interactions. Studies following this approach have shown evidence for both competition and independence between SL and EF (Horvath et al., 2023; Vékony et al., 2019).

While most of these studies focus on the initial acquisition of probabilistic knowledge, their methodological approaches can also be extended to study the relationship between rewiring and EF. Our study follows the first approach, whereas Horvath et al. (2022) follows the fourth. In Horvath et al., participants actively inhibited motor responses associated with the knowledge acquired in the Learning Phase, whereas in our study, we measured interindividual differences in inhibitory control capacity. This methodological difference may explain the discrepancy in results. It is possible that, in Horvath et al., individuals’ cognitive inhibitory resources were allocated to refraining from specific motor responses rather than inhibiting implicit probabilistic representations acquired in the Learning Phase. These results can be interpreted within a dual-task framework, which suggests that if a secondary task impairs performance, the cognitive function engaged in the secondary task is crucial for achieving the good performance in the primary task (Fisk et al., 1986). Thus, in Horvath et al. (2022), the weaker rewiring scores observed for inhibited (no-go) trials compared to go trials may indicate that inhibitory capacity is essential for the rewiring process itself.

Conversely to the positive relationship between rewiring and inhibition observed in our study, we found a negative correlation between rewiring and semantic fluency. This result suggests that individuals with higher semantic fluency may be less proficient at updating their implicit probabilistic representations in response to environmental changes. While this result might seem unexpected within the broader context of EF and habit change research (Allom et al., 2018), it is consistent with the competitive learning and memory systems framework (Ambrus et al., 2020; Janacsek et al., 2012; Lee et al., 2014; Pedraza et al., 2024; Smittenaar et al., 2013). Given that verbal fluency tasks, especially semantic fluency, involves the controlled access to long term memory representations (Kavé & Sapir-Yogev, 2020; Marko et al., 2023; Ullman, 2013), a possible interpretation for our result could be that individuals with greater verbal fluency rely more heavily on pre-existing cognitive mnesic models, which could impede their ability to integrate novel probabilistic information (Pedraza et al., 2024). This negative relationship aligns with the competition hypothesis of dual-process neuro-computational models where a greater reliance on model-based cognitive functioning is associated with a smaller reliance on model-free processes (Daw et al., 2005; Janacsek et al., 2012; Nemeth, Janacsek, & Fiser, 2013; Poldrack & Packard, 2003). In this context, proficiency in semantic fluency could reflect more efficient access to long-term memory, suggesting a heavier reliance on model-based processes. This reliance may hinder the acquisition of new probabilistic information, which may be dependent on the model-free system. Similar results were also reported by Pedraza et al. (2024), suggesting that the ability to acquire new probabilistic representations might be related to a reduced reliance on model-based functioning, as reflected by poorer performance on verbal fluency tasks.

It is important to note, however, that while verbal fluency tasks are generally considered good indicators of EF (Amunts et al., 2020; Gustavson et al., 2019), their precise role within the Unity and Diversity model of EF (Miyake et al., 2000) is still debated. Studies examining verbal fluency tasks in the context of Miyake’s model have suggested that these tasks may reflect cognitive flexibility or set shifting (Gustavson et al., 2019; Troyer et al., 1997). Based on this, it could be hypothesized that the negative correlation between rewiring and verbal fluency found in our study reflects competition between updating probabilistic representations and cognitive flexibility. Nonetheless, this interpretation seems unlikely given our data, as no relationship was found between rewiring and BCST, a well-established set-shifting task (Piper et al., 2015). Additionally, the lack of correlation between BCST and verbal fluency tasks in our analysis further suggests that verbal fluency tasks are not purely measures of cognitive flexibility. Semantic, lexical, and action fluency tasks likely involve different cognitive processes and may differ to what extent they reflect EF. For example, while cognitive flexibility has been shown to be the primary factor in the lexical fluency task, the semantic fluency task may also involve other components such as categorical production (Troyer et al., 1997), which likely reflects long-term memory access (Freedman & Loftus, 1971). These distinctions support our interpretation that semantic fluency reflects model-based functioning, particularly within the context of the competition hypothesis of dual-process neuro-computational processes.

Verbal fluency tasks have also been argued to reflect working memory updating (Azuma, 2004; Daneman, 1991; Shao et al., 2014). In the context of our results, this could imply that working memory updating proficiency might counteract the rewiring of probabilistic representations. However, this interpretation seems unlikely given our data, as we observed no relationship between the Counting Span task, a valid and reliable working memory task (Conway et al., 2005), and rewiring or verbal fluency. In our previous study (Pedraza et al., 2024), using exploratory factor analysis we have identified a common factor underlying both the Counting Span and verbal fluency tasks, which negatively correlated with SL. We have suggested that this factor could reflect either updating or access to long-term memory models. Based on our current results, the latter explanation seems more plausible. This interpretation aligns with Kurdi et al. (2019), who suggest that the updating of implicit probabilistic representations depends exclusively on model-free processes. It is also important to note that the updating referenced in the context of EF models differs from the updating involved in the probabilistic learning and neuro-computational modeling framework. The former involves the conscious maintenance and manipulation of information in working memory (Ecker et al., 2014; Miyake & Friedman, 2012), whereas the latter involves the implicit modification of informational priors (Friston, 2010; Pesthy et al., 2023). This is consistent with our interpretation that not the updating in working memory, but the access to long-term memory affects rewiring performance.

Our correlational analyses results were further corroborated through multiple regression analysis, which revealed that inhibition, measured by the GoNoGo task, positively predicted rewiring performance, while semantic fluency negatively predicted rewiring it. This statistical approach strengthened the validity of our results and provided additional support for the observed relationships between executive functions and the rewiring of probabilistic representations (Crawford, 2006). Our findings highlight the complex cognitive dynamics involved in the rewiring of implicit probabilistic representations. Successful rewiring appears to require a balance between inhibiting outdated cognitive schemas and acquiring new probabilistic information. Different cognitive functions, such as inhibition and verbal fluency, may contribute to these processes in distinct ways. Inhibition may facilitate the suppression of motor responses and memory representations related to outdated knowledge, facilitating the integration of new information (Danner et al., 2011; Jahanshahi et al., 2015). Conversely, verbal fluency may reflect a greater reliance on model-based functioning, which could hinder the flexibility needed to acquire novel probabilistic representations (Janacsek et al., 2012; Kavé & Sapir-Yogev, 2020; Marko et al., 2023; Pedraza et al., 2024).

These findings carry significant theoretical implications for understanding the cognitive mechanisms involved in acquiring probabilistic information across various contexts. In the literature, there has been ongoing discourse regarding whether the acquisition of new information, whether under conditions of proactive interference or not, relies on similar or distinct neural and cognitive mechanisms (Berman & Dudai, 2001; Bouton, 2000; Horváth et al., 2022). The negative correlation we observed between semantic fluency and rewiring performance echoes findings from studies investigating the link between EF and learning (Pedraza et al., 2024; Virag et al., 2015). This suggests a possible convergence in the neurocognitive mechanisms that support both learning and rewiring processes. Nonetheless, this hypothesis warrants further investigation, particularly through studies exploring the cellular and neural architectural dynamics underlying learning and rewiring of probabilistic information.

Our findings also highlight the potential for cognitive control interventions, particularly those targeting inhibitory control, to enhance the rewiring process. Research in fields such as food intake and addiction has shown that cognitive control training can effectively modify habitual behaviors (Allom et al., 2018; Galla & Duckworth, 2015; Preuss et al., 2017). Bolstering inhibitory control could enable individuals to better suppress outdated cognitive models and more easily integrate new information, facilitating adaptive learning and habit updating. Nonetheless, it is important to acknowledge the limitations of this study. For instance, the Rewiring Phase and executive function measurements were conducted on separate days. Intraindividual differences in the reliance on model-based or model-free functioning may be influenced by factors such as emotional state (Sherman et al., 2024; Tóth-Fáber et al., 2020) or hormonal fluctuations during the menstrual cycle (Diekhof et al., 2021; Joue et al., 2022). These factors could contribute to variability in our findings. Future studies should address these limitations to improve the robustness and generalizability of the results. Additionally, incorporating neuroimaging techniques such as fMRI or EEG could elucidate the neural mechanisms underlying the interplay between these cognitive processes.

To our knowledge, this study is the first to explore the relationship between the rewiring of implicit probabilistic information and a wide range of prefrontal-dependent executive functions. Our findings suggests a nuanced interaction between these cognitive processes, as revealed through the study of individual differences: while rewiring showed a positive relationship with cognitive inhibition, it was negatively associated with verbal fluency. This indicates that the rewiring of implicit probabilistic representations is a multifaceted cognitive process intricately intertwined with executive functions. It appears to necessitate both overcoming proactive interference from prior knowledge, facilitated by cognitive inhibition, and a reliance on model-free functioning to integrate novel probabilistic information, as evidenced by the negative relationship with semantic fluency. By shedding light on the relationship between executive functions and rewiring, this study provides a foundational understanding of the cognitive dynamics underlying habit change.

## Acknowledgments

The authors are grateful to Jeoffrey Maillard for managing participant enrollment. This work was supported by the Chaire de Professeur Junior Program by INSERM and the French National Grant Agency (Chaire Inserm-ANR-22-CE64-001).

## Data availability statement

In accordance with the transparency and reproducibility principles, we provide information regarding the availability of data supporting the findings of this study. Code and data necessary to replicate the findings of this manuscript can be found on the OSF platform, at the following link:

https://osf.io/gqpjn/

## Declaration of generative AI and AI-assisted technologies in the writing process

During the preparation of this work the authors used ChatGPT-4o in order to improve readability and language. After using this tool, the authors thoroughly reviewed and edited the content as needed and take full responsibility for the content of the published article.

## Supplementary material

EF_and_rewiring_supl

## References

Allan, J. L., McMinn, D., & Daly, M. (2016). A Bidirectional Relationship between Executive Function and Health Behavior: Evidence, Implications, and Future Directions. Frontiers in Neuroscience, 10. 10.3389/fnins.2016.00386

Allom, V., Mullan, B., & Hagger, M. (2016). Does inhibitory control training improve health behaviour? A meta-analysis. Health Psychology Review, 10(2), 168–186. 10.1080/17437199.2015.1051078

Allom, V., Mullan, B., Smith, E., Hay, P., & Raman, J. (2018). Breaking bad habits by improving executive function in individuals with obesity. BMC Public Health, 18(1), 505. 10.1186/s12889-018-5392-y

Ambrus, G. G., Vékony, T., Janacsek, K., Trimborn, A. B. C., Kovács, G., & Nemeth, D. (2020). When less is more: Enhanced statistical learning of non-adjacent dependencies after disruption of bilateral DLPFC. Journal of Memory and Language, 114, 104144. 10.1016/j.jml.2020.104144

Amunts, J., Camilleri, J. A., Eickhoff, S. B., Heim, S., & Weis, S. (2020). Executive functions predict verbal fluency scores in healthy participants. Scientific Reports, 10(1), 11141. 10.1038/s41598-020-65525-9

Aslin, R. N. (2017). Statistical learning: A powerful mechanism that operates by mere exposure. WIREs Cognitive Science, 8(1–2), e1373. 10.1002/wcs.1373

Azuma, T. (2004). Working Memory and Perseveration in Verbal Fluency. Neuropsychology, 18, 69–77. 10.1037/0894-4105.18.1.69

Badre, D., Kayser, A. S., & D’Esposito, M. (2010). Frontal Cortex and the Discovery of Abstract Action Rules. Neuron, 66(2), 315–326. 10.1016/j.neuron.2010.03.025

Baldwin, D., Andersson, A., Saffran, J., & Meyer, M. (2008). Segmenting dynamic human action via statistical structure. Cognition, 106(3), 1382–1407. 10./j.cognition.2007.07.005

Beierholm, U. R., Anen, C., Quartz, S., & Bossaerts, P. (2011). Separate encoding of model-based and model-free valuations in the human brain. NeuroImage, 58(3), 955–962. 10.1016/j.neuroimage.2011.06.071

Berg, E. A. (1948). A Simple Objective Technique for Measuring Flexibility in Thinking. The Journal of General Psychology. https://www.tandfonline.com/doi/abs/10.1080/00221309.1948.9918159

Berman, D. E., & Dudai, Y. (2001). Memory Extinction, Learning Anew, and Learning the New: Dissociations in the Molecular Machinery of Learning in Cortex. Science, 291(5512), 2417–2419. 10.1126/science.1058165

Borragán, G., Slama, H., Destrebecqz, A., & Peigneux, P. (2016). Cognitive Fatigue Facilitates Procedural Sequence Learning. Frontiers in Human Neuroscience, 10. 10.3389/fnhum.2016.00086

Bouton, M. E. (2000). A learning theory perspective on lapse, relapse, and the maintenance of behavior change. Health Psychology, 19(1S), 57–63. 10.1037/0278-6133.19.suppl1.57

Braver, T. S., Reynolds, J. R., & Donaldson, D. I. (2003). Neural Mechanisms of Transient and Sustained Cognitive Control during Task Switching. Neuron, 39(4), 713–726. 10.1016/S0896-6273(03)00466-5

Buffington, J., Demos, A. P., & Morgan-Short, K. (2021). The Reliability and Validity of Procedural Memory Assessments Used in Second Language Acquisition Research. Studies in Second Language Acquisition, 43(3), 635–662. 10.1017/S0272263121000127

Carden, L., & Wood, W. (2018). Habit formation and change. Current Opinion in Behavioral Sciences, 20, 117–122. 10.1016/j.cobeha.2017.12.009

Case, R., Kurland, D. M., & Goldberg, J. (1982). Operational efficiency and the growth of short-term memory span. Journal of Experimental Child Psychology, 33(3), 386–404. 10.1016/0022-0965(82)90054-6

Conway, A. R. A., Kane, M. J., Bunting, M. F., Hambrick, D. Z., Wilhelm, O., & Engle, R. W. (2005). Working memory span tasks: A methodological review and user’s guide. Psychonomic Bulletin & Review, 12(5), 769–786. 10.3758/BF03196772

Conway, C. M. (2020). How does the brain learn environmental structure? Ten core principles for understanding the neurocognitive mechanisms of statistical learning. Neuroscience and Biobehavioral Reviews, 112, 279–299. 10.1016/j.neubiorev.2020.01.032

Costa, K. G., Cabral, D. A., Hohl, R., & Fontes, E. B. (2019). Rewiring the Addicted Brain Through a Psychobiological Model of Physical Exercise. Frontiers in Psychiatry, 10. 10.3389/fpsyt.2019.00600

Crawford, S. L. (2006). Correlation and Regression. Circulation, 114(19), 2083–2088. 10.1161/CIRCULATIONAHA.105.586495

Curtis, C. E., & D’Esposito, M. (2003). Persistent activity in the prefrontal cortex during working memory. Trends in Cognitive Sciences, 7(9), 415–423. 10.1016/S1364-6613(03)00197-9

Daneman, M. (1991). Working memory as a predictor of verbal fluency. Journal of Psycholinguistic Research, 20(6), 445–464. 10.1007/BF01067637

Danner, U. N., Aarts, H., Papies, E. K., & de Vries, N. K. (2011). Paving the path for habit change: Cognitive shielding of intentions against habit intrusion. British Journal of Health Psychology, 16(1), 189–200. 10.1348/2044-8287.002005

Daw, N. D., Niv, Y., & Dayan, P. (2005). Uncertainty-based competition between prefrontal and dorsolateral striatal systems for behavioral control. Nature Neuroscience, 8(12), 1704–1711. 10.1038/nn1560

Decker, J. H., Otto, A. R., Daw, N. D., & Hartley, C. A. (2016). From Creatures of Habit to Goal-Directed Learners: Tracking the Developmental Emergence of Model-Based Reinforcement Learning. Psychological Science, 27(6), 848–858. 10.1177/0956797616639301

Diamond, A. (2013). Executive Functions. Annual Review of Psychology, 64(1), 135–168. 10.1146/annurev-psych-113011-143750

Dickinson, A., & Balleine, B. (1994). Motivational control of goal-directed action. Animal Learning & Behavior, 22(1), 1–18. 10.3758/BF03199951

Dickinson, A., & Weiskrantz, L. (1997). Actions and habits: The development of behavioural autonomy. Philosophical Transactions of the Royal Society of London. B, Biological Sciences, 308(1135), 67–78. 10.1098/rstb.1985.0010

Diekhof, E. K., Geana, A., Ohm, F., Doll, B. B., & Frank, M. J. (2021). The Straw That Broke the Camel’s Back: Natural Variations in 17β-Estradiol and COMT-Val158Met Genotype Interact in the Modulation of Model-Free and Model-Based Control. Frontiers in Behavioral Neuroscience, 15. 10.3389/fnbeh.2021.658769

Dienes, Z. (2014). Using Bayes to get the most out of non-significant results. Frontiers in Psychology, 5. 10.3389/fpsyg.2014.00781

Doll, B. B., Shohamy, D., & Daw, N. D. (2015). Multiple memory systems as substrates for multiple decision systems. Neurobiology of Learning and Memory, 117, 4–13. 10.1016/j.nlm.2014.04.014

Doyon, J., Bellec, P., Amsel, R., Penhune, V., Monchi, O., Carrier, J., Lehéricy, S., & Benali, H. (2009). Contributions of the basal ganglia and functionally related brain structures to motor learning. Behavioural Brain Research, 199(1), 61–75. 10.1016/j.bbr.2008.11.012

Dresp-Langley, B. (2012). Why the Brain Knows More than We Do: Non-Conscious Representations and Their Role in the Construction of Conscious Experience. Brain Sciences, 2(1), Article 1. 10.3390/brainsci2010001

Ecker, U. K. H., Lewandowsky, S., & Oberauer, K. (2014). Removal of information from working memory: A specific updating process. Journal of Memory and Language, 74, 77–90. 10.1016/j.jml.2013.09.003

Fan, J., McCandliss, B. D., Fossella, J., Flombaum, J. I., & Posner, M. I. (2005). The activation of attentional networks. NeuroImage, 26(2), 471–479. 10.1016/j.neuroimage.2005.02.004

Fan, J., McCandliss, B. D., Sommer, T., Raz, A., & Posner, M. I. (2002). Testing the Efficiency and Independence of Attentional Networks. Journal of Cognitive Neuroscience, 14(3), 340–347. 10.1162/089892902317361886

Farkas, B. C., Krajcsi, A., Janacsek, K., & Nemeth, D. (2024). The complexity of measuring reliability in learning tasks: An illustration using the Alternating Serial Reaction Time Task. Behavior Research Methods, 56(1), 301–317. 10.3758/s13428-022-02038-5

Filoteo, J. V., Lauritzen, S., & Maddox, W. T. (2010). Removing the frontal lobes: The effects of engaging executive functions on perceptual category learning. Psychological Science, 21(3), 415–423. 10.1177/0956797610362646

Fisk, A. D., Derrick, W. L., & Schneider, W. (1986). A methodological assessment and evaluation of dual-task paradigms. Current Psychological Research & Reviews, 5(4), 315–327. 10.1007/BF02686599

Foerde, K. (2018). What are habits and do they depend on the striatum? A view from the study of neuropsychological populations. Current Opinion in Behavioral Sciences, 20, 17–24. 10.1016/j.cobeha.2017.08.011

Fox, C. J., Mueller, S. T., Gray, H. M., Raber, J., & Piper, B. J. (2013). Evaluation of a Short-Form of the Berg Card Sorting Test. PLOS ONE, 8(5), e63885. 10.1371/journal.pone.0063885

Freedman, J. L., & Loftus, E. F. (1971). Retrieval of words from long-term memory. Journal of Verbal Learning and Verbal Behavior, 10(2), 107–115. 10.1016/S0022-5371(71)80001-4

Friston, K. (2010). The free-energy principle: A unified brain theory? Nature Reviews Neuroscience, 11(2), 127–138. 10.1038/nrn2787

Funahashi, S., & Andreau, J. M. (2013). Prefrontal cortex and neural mechanisms of executive function. Journal of Physiology-Paris, 107(6), 471–482. 10.1016/j.jphysparis.2013.05.001

Galla, B. M., & Duckworth, A. L. (2015). More than Resisting Temptation: Beneficial Habits Mediate the Relationship between Self-Control and Positive Life Outcomes. Journal of Personality and Social Psychology, 109(3), 508–525. 10.1037/pspp0000026

Goedert, K. M., & Willingham, D. B. (2002). Patterns of Interference in Sequence Learning and Prism Adaptation Inconsistent With the Consolidation Hypothesis. Learning & Memory, 9(5), 279–292. 10.1101/lm.50102

Graybiel, A. M. (2008). Habits, Rituals, and the Evaluative Brain. Annual Review of Neuroscience, 31(Volume 31, 2008), 359–387. 10.1146/annurev.neuro.29.051605.112851

Gray-Burrows, K., Taylor, N., O’Connor, D., Sutherland, E., Stoet, G., & Conner, M. (2019). A systematic review and meta-analysis of the executive function-health behaviour relationship. Health Psychology and Behavioral Medicine, 7(1), 253–268. 10.1080/21642850.2019.1637740

Greenwald, A. G., & Banaji, M. R. (1995). Implicit social cognition: Attitudes, self-esteem, and stereotypes. Psychological Review, 102(1), 4–27. 10.1037/0033-295X.102.1.4

Gustavson, D. E., Panizzon, M. S., Franz, C. E., Reynolds, C. A., Corley, R. P., Hewitt, J. K., Lyons, M. J., Kremen, W. S., & Friedman, N. P. (2019). Integrating verbal fluency with executive functions: Evidence from twin studies in adolescence and middle age. Journal of Experimental Psychology: General, 148, 2104–2119. 10.1037/xge0000589

Hagger, M. S., Cameron, L. D., Hamilton, K., Hankonen, N., & Lintunen, T. (2020). The Handbook of Behavior Change. Cambridge University Press.

Hallgató, E., Győri-Dani, D., Pekár, J., Janacsek, K., & Nemeth, D. (2013). The differential consolidation of perceptual and motor learning in skill acquisition. Cortex, 49(4), 1073–1081. 10.1016/j.cortex.2012.01.002

Harrison, J. E., Buxton, P., Husain, M., & Wise, R. (2000). Short test of semantic and phonological fluency: Normal performance, validity and test-retest reliability. British Journal of Clinical Psychology, 39(2), 181–191. 10.1348/014466500163202

Hartley, T., & Burgess, N. (2005). Complementary memory systems: Competition, cooperation and compensation. Trends in Neurosciences, 28(4), 169–170. 10.1016/j.tins.2005.02.004

Henke, K. (2010). A model for memory systems based on processing modes rather than consciousness. Nature Reviews. Neuroscience, 11(7), 523–532. 10.1038/nrn2850

Hong, I., Kim, M.-S., & Jeong, S. K. (2022). Flexibility and stability of habit learning depend on temporal signal variation. Journal of Experimental Psychology: Learning, Memory, and Cognition, 48(1), 1–12. 10.1037/xlm0001113

Horvath, K., Nemeth, D., Janacsek, K., & Kóbor, A. (2023). Independent and interactive dynamics between statistical learning and inhibitory control. 10.31234/osf.io/ptdkr

Horváth, K., Nemeth, D., & Janacsek, K. (2022). Inhibitory control hinders habit change. Scientific Reports, 12(1), Article 1. 10.1038/s41598-022-11971-6

Howard, J. H., & Howard, D. V. (1997). Age differences in implicit learning of higher order dependencies in serial patterns. Psychology and Aging, 12(4), 634–656.

Isbilen, E. S., & Christiansen, M. H. (2022). Statistical learning of language: A metaC:analysis into 25 years of research. Cognitive Science, 46(9), e13198.

Jahanshahi, M., Obeso, I., Rothwell, J. C., & Obeso, J. A. (2015). A fronto–striato– subthalamic–pallidal network for goal-directed and habitual inhibition. Nature Reviews Neuroscience, 16(12), 719–732. 10.1038/nrn4038

Janacsek, K., Fiser, J., & Nemeth, D. (2012). The best time to acquire new skills: Age-related differences in implicit sequence learning across the human lifespan. Developmental Science, 15(4), 496–505. 10.1111/j.1467-7687.2012.01150.x

Janacsek, K., & Nemeth, D. (2012). Predicting the future: From implicit learning to consolidation. International Journal of Psychophysiology, 83(2), 213–221. 10.1016/j.ijpsycho.2011.11.012

Jiang, Y. V., & Sisk, C. A. (2019). Habit-like attention. Current Opinion in Psychology, 29, 65–70. 10.1016/j.copsyc.2018.11.014

Joue, G., Chakroun, K., Bayer, J., Gläscher, J., Zhang, L., Fuss, J., Hennies, N., & Sommer, T. (2022). Sex Differences and Exogenous Estrogen Influence Learning and Brain Responses to Prediction Errors. Cerebral Cortex, 32(9), 2022–2036. 10.1093/cercor/bhab334

Kaufman, S. B., Deyoung, C. G., Gray, J. R., Jiménez, L., Brown, J., & Mackintosh, N. (2010). Implicit learning as an ability. Cognition, 116(3), 321–340. 10.1016/j.cognition.2010.05.011

Kavé, G., & Sapir-Yogev, S. (2020). Associations between memory and verbal fluency tasks. Journal of Communication Disorders, 83, 105968. 10.1016/j.jcomdis.2019.105968

Kóbor, A., Janacsek, K., Takács, Á., & Nemeth, D. (2017). Statistical learning leads to persistent memory: Evidence for one-year consolidation. Scientific Reports, 7(1), 760. 10.1038/s41598-017-00807-3

Koechlin, E. (2016). Prefrontal executive function and adaptive behavior in complex environments. Current Opinion in Neurobiology, 37, 1–6. 10.1016/j.conb.2015.11.004

Kondo, H., Osaka, N., & Osaka, M. (2004). Cooperation of the anterior cingulate cortex and dorsolateral prefrontal cortex for attention shifting. NeuroImage, 23(2), 670–679. 10.1016/j.neuroimage.2004.06.014

Kurdi, B., Gershman, S. J., & Banaji, M. R. (2019). Model-free and model-based learning processes in the updating of explicit and implicit evaluations. Proceedings of the National Academy of Sciences, 116(13), 6035–6044. 10.1073/pnas.1820238116

Langner, R., Scharnowski, F., Ionta, S., Salmon, C. E. G., Piper, B. J., & Pamplona, G. S. P. (2023). Evaluation of the reliability and validity of computerized tests of attention. PLOS ONE, 18(1), e0281196. 10.1371/journal.pone.0281196

Lee, S. W., Shimojo, S., & O’Doherty, J. P. (2014). Neural Computations Underlying Arbitration between Model-Based and Model-free Learning. Neuron, 81(3), 687–699. 10.1016/j.neuron.2013.11.028

Logan, G. D. (1988). Toward an instance theory of automatization. Psychological Review, 95(4), 492–527. 10.1037/0033-295X.95.4.492

Marko, M., Michalko, D., Dragašek, J., Vančová, Z., Jarčušková, D., & Riečanský, I. (2023). Assessment of Automatic and Controlled Retrieval Using Verbal Fluency Tasks. Assessment, 30(7), 2198–2211. 10.1177/10731911221117512

Miller, E. K., & Cohen, J. D. (2001). An integrative theory of prefrontal cortex function. Annual Review of Neuroscience, 24, 167–202. 10.1146/annurev.neuro.24.1.167

Miyake, A., & Friedman, N. P. (2012). The Nature and Organization of Individual Differences in Executive Functions: Four General Conclusions. Current Directions in Psychological Science, 21(1), 8–14. 10.1177/0963721411429458

Miyake, A., Friedman, N. P., Emerson, M. J., Witzki, A. H., Howerter, A., & Wager, T. D. (2000). The unity and diversity of executive functions and their contributions to complex “Frontal Lobe” tasks: A latent variable analysis. Cognitive Psychology, 41(1), 49–100. 10.1006/cogp.1999.0734

Mueller, S. T., & Piper, B. J. (2014a). The Psychology Experiment Building Language (PEBL) and PEBL Test Battery. Journal of Neuroscience Methods, 222, 250–259. 10.1016/j.jneumeth.2013.10.024

Mueller, S. T., & Piper, B. J. (2014b). The Psychology Experiment Building Language (PEBL) and PEBL Test Battery. Journal of Neuroscience Methods, 222, 250–259. 10.1016/j.jneumeth.2013.10.024

Nemeth, D., Janacsek, K., & Fiser, J. (2013). Age-dependent and coordinated shift in performance between implicit and explicit skill learning. Frontiers in Computational Neuroscience, 7. https://www.frontiersin.org/articles/10.3389/fncom.2013.00147

Nemeth, D., Janacsek, K., Polner, B., & Kovacs, Z. A. (2013). Boosting Human Learning by Hypnosis. Cerebral Cortex, 23(4), 801–805. 10.1093/cercor/bhs068

Osaka, M., Osaka, N., Kondo, H., Morishita, M., Fukuyama, H., Aso, T., & Shibasaki, H. (2003). The neural basis of individual differences in working memory capacity: An fMRI study. NeuroImage, 18(3), 789–797. 10.1016/S1053-8119(02)00032-0

Osaka, N., Osaka, M., Kondo, H., Morishita, M., Fukuyama, H., & Shibasaki, H. (2004). The neural basis of executive function in working memory: An fMRI study based on individual differences. NeuroImage, 21(2), 623–631. 10.1016/j.neuroimage.2003.09.069

Otto, A. R., Skatova, A., Madlon-Kay, S., & Daw, N. D. (2015). Cognitive Control Predicts Use of Model-based Reinforcement Learning. Journal of Cognitive Neuroscience, 27(2), 319–333. 10.1162/jocn_a_00709

Packard, M. G., Hirsh, R., & White, N. M. (1989). Differential effects of fornix and caudate nucleus lesions on two radial maze tasks: Evidence for multiple memory systems. The Journal of Neuroscience: The Official Journal of the Society for Neuroscience, 9(5), 1465–1472.

Palminteri, S., Kilford, E. J., Coricelli, G., & Blakemore, S.-J. (2016). The Computational Development of Reinforcement Learning during Adolescence. PLOS Computational Biology, 12(6), e1004953. 10.1371/journal.pcbi.1004953

Paneri, S., & Gregoriou, G. G. (2017). Top-Down Control of Visual Attention by the Prefrontal Cortex. Functional Specialization and Long-Range Interactions. Frontiers in Neuroscience, 11. 10.3389/fnins.2017.00545

Park, J., Janacsek, K., Nemeth, D., & Jeon, H.-A. (2022). Reduced functional connectivity supports statistical learning of temporally distributed regularities. NeuroImage, 260, 119459. 10.1016/j.neuroimage.2022.119459

Park, J., Yoon, H.-D., Yoo, T., Shin, M., & Jeon, H.-A. (2020). Potential and efficiency of statistical learning closely intertwined with individuals’ executive functions: A mathematical modeling study. Scientific Reports, 10(1), Article 1. 10.1038/s41598-020-75157-8

Parks, K. M. A., Griffith, L. A., Armstrong, N. B., & Stevenson, R. A. (2020). Statistical Learning and Social Competency: The Mediating Role of Language. Scientific Reports, 10(1), Article 1. 10.1038/s41598-020-61047-6

Pedraza, F., Farkas, B. C., Vékony, T., Haesebaert, F., Phelipon, R., Mihalecz, I., Janacsek, K., Anders, R., Tillmann, B., Plancher, G., & Németh, D. (2024). Evidence for a competitive relationship between executive functions and statistical learning. Npj Science of Learning, 9(1), 1–14. 10.1038/s41539-024-00243-9

Pesthy, O., Farkas, K., Sapey-Triomphe, L.-A., Guttengéber, A., Komoróczy, E., Janacsek, K., Réthelyi, J. M., & Németh, D. (2023). Intact predictive processing in autistic adults: Evidence from statistical learning. Scientific Reports, 13(1), 11873. 10.1038/s41598-023-38708-3

Peretz, I., Saffran, J., Schön, D., & Gosselin, N. (2012). Statistical learning of speech, not music, in congenital amusia. Annals of the New York Academy of Sciences, 1252, 361–367. 10.1111/j.1749-6632.2011.06429.x

Phelps, E. A., Hyder, F., Blamire, A. M., & Shulman, R. G. (1997). FMRI of the prefrontal cortex during overt verbal fluency. NeuroReport, 8(2), 561.

Piper, B. J., Mueller, S. T., Geerken, A. R., Dixon, K. L., Kroliczak, G., Olsen, R. H. J., & Miller, J. K. (2015). Reliability and validity of neurobehavioral function on the Psychology Experimental Building Language test battery in young adults. PeerJ, 3, e1460. 10.7717/peerj.1460

Poldrack, R. A., & Packard, M. G. (2003). Competition among multiple memory systems: Converging evidence from animal and human brain studies. Neuropsychologia, 41(3), 245–251.

Poldrack, R. A. (2022). Hard to Break: Why Our Brains Make Habits Stick. Princeton University Press.

Preuss, H., Pinnow, M., Schnicker, K., & Legenbauer, T. (2017). Improving Inhibitory Control Abilities (ImpulsE)—A Promising Approach to Treat Impulsive Eating? European Eating Disorders Review, 25(6), 533–543. 10.1002/erv.2544

Robinson, G., Shallice, T., Bozzali, M., & Cipolotti, L. (2012). The differing roles of the frontal cortex in fluency tests. Brain, 135(7), 2202–2214. 10.1093/brain/aws142

Rohrmeier, M., & Rebuschat, P. (2012). Implicit Learning and Acquisition of Music. Topics in Cognitive Science, 4(4), 525–553. 10.1111/j.1756-8765.2012.01223.x

Romano Bergstrom, J. C., Howard, J. H., & Howard, D. V. (2012). Enhanced implicit sequence learning in college-age video game players and musicians. Applied Cognitive Psychology, 26(1), 91–96. 10.1002/acp.1800

Rossi, A. F., Pessoa, L., Desimone, R., & Ungerleider, L. G. (2009). The prefrontal cortex and the executive control of attention. Experimental Brain Research, 192(3), 489–497. 10.1007/s00221-008-1642-z

Ruffman, T., Taumoepeau, M., & Perkins, C. (2012). Statistical learning as a basis for social understanding in children. The British Journal of Developmental Psychology, 30(Pt 1), 87–104. 10.1111/j.2044-835X.2011.02045.x

Saffran, J. R., Aslin, R. N., & Newport, E. L. (1996). Statistical Learning by 8-Month-Old Infants. Science, 274(5294), 1926–1928. 10.1126/science.274.5294.1926

Seger, C. A., & Spiering, B. J. (2011). A Critical Review of Habit Learning and the Basal Ganglia. Frontiers in Systems Neuroscience, 5. 10.3389/fnsys.2011.00066

Shao, Z., Janse, E., Visser, K., & Meyer, A. S. (2014). What do verbal fluency tasks measure? Predictors of verbal fluency performance in older adults. Frontiers in Psychology, 5, 772. 10.3389/fpsyg.2014.00772

Sherman, B. E., Huang, I., Wijaya, E. G., Turk-Browne, N. B., & Goldfarb, E. V. (2024). Acute Stress Effects on Statistical Learning and Episodic Memory. Journal of Cognitive Neuroscience, 36(8), 1741–1759. 10.1162/jocn_a_02178

Shohamy, D., & Daw, N. D. (2014). Habits and reinforcement learning. In The cognitive neurosciences, 5th ed (pp. 591–603). MIT Press.

Smalle, E. H. M., Daikoku, T., Szmalec, A., Duyck, W., & Möttönen, R. (2022). Unlocking adults’ implicit statistical learning by cognitive depletion. Proceedings of the National Academy of Sciences, 119(2), e2026011119. 10.1073/pnas.2026011119

Smittenaar, P., FitzGerald, T. H. B., Romei, V., Wright, N. D., & Dolan, R. J. (2013). Disruption of dorsolateral prefrontal cortex decreases model-based in favor of model-free control in humans. Neuron, 80(4), Article 4. 10.1016/j.neuron.2013.08.009

Song, S., Howard, J. H., & Howard, D. V. (2007). Implicit probabilistic sequence learning is independent of explicit awareness. Learning & Memory, 14(3), 167–176. 10.1101/lm.437407

St-Hilaire, A., Hudon, C., Vallet, G. T., Bherer, L., Lussier, M., Gagnon, J.-F., Simard, M., Gosselin, N., Escudier, F., Rouleau, I., & Macoir, J. (2016). Normative data for phonemic and semantic verbal fluency test in the adult French-Quebec population and validation study in Alzheimer’s disease and depression. The Clinical Neuropsychologist, 30(7), 1126–1150. 10.1080/13854046.2016.1195014

Szegedi-Hallgató, E., Janacsek, K., Vékony, T., Tasi, L. A., Kerepes, L., Hompoth, E. A., Bálint, A., & Németh, D. (2017). Explicit instructions and consolidation promote rewiring of automatic behaviors in the human mind. Scientific Reports, 7(1), Article 1. 10.1038/s41598-017-04500-3

Thompson, C., & Sabik, M. (2018). Allocation of attention in familiar and unfamiliar traffic scenarios. Transportation Research Part F: Traffic Psychology and Behaviour, 55, 188–198. 10.1016/j.trf.2018.03.006

Tillmann, B., & McAdams, S. (2004). Implicit Learning of Musical Timbre Sequences: Statistical Regularities Confronted With Acoustical (Dis)Similarities. Journal of Experimental Psychology: Learning, Memory, and Cognition, 30(5), 1131–1142. 10.1037/0278-7393.30.5.1131

Tillmann, B., & Poulin-Charronnat, B. (2010). Auditory expectations for newly acquired structures. Quarterly Journal of Experimental Psychology, 63(8), 1646–1664.

Tóth, B., Janacsek, K., Takács, Á., Kóbor, A., Zavecz, Z., & Nemeth, D. (2017). Dynamics of EEG functional connectivity during statistical learning. Neurobiology of Learning and Memory, 144, 216–229. 10.1016/j.nlm.2017.07.015

Tóth-Fáber, E., Janacsek, K., Szőllősi, Á., Kéri, S., & Németh, D. (2020). Regularity extraction under stress: Boosted statistical learning but unaffected sequence learning (p. 2020.05.13.092726). bioRxiv. 10.1101/2020.05.13.092726

Troyer, A. K., Moscovitch, M., & Winocur, G. (1997). Clustering and switching as two components of verbal fluency: Evidence from younger and older healthy adults. Neuropsychology, 11(1), 138–146. 10.1037//0894-4105.11.1.138

Tuhkanen, S., Pekkanen, J., Rinkkala, P., Mole, C., Wilkie, R. M., & Lappi, O. (2019). Humans Use Predictive Gaze Strategies to Target Waypoints for Steering. Scientific Reports, 9(1), 8344. 10.1038/s41598-019-44723-0

Ullman, M. T. (2001). The declarative/procedural model of lexicon and grammar. Journal of psycholinguistic research, 30, 37–69.

Ullman, M. T. (2013). The role of declarative and procedural memory in disorders of language. Linguistic Variation, 13(2), 133–154. 10.1075/lv.13.2.01ull

Ullman, M. T. (2020). The declarative/procedural model: A neurobiologically motivated theory of first and second language 1. In Theories in second language acquisition (pp. 128–161). Routledge.

Vékony, T., Ambrus, G. G., Janacsek, K., & Nemeth, D. (2022). Cautious or causal? Key implicit sequence learning paradigms should not be overlooked when assessing the role of DLPFC (Commentary on Prutean et al.). Cortex, 148, 222–226. 10.1016/j.cortex.2021.10.001

Vékony, T., Török, L., Pedraza, F., Schipper, K., Plèche, C., Tóth, L., Janacsek, K., & Nemeth, D. (2019). Retrieval of a well-established skill is resistant to distraction: Evidence from an implicit probabilistic sequence learning task. bioRxiv, 849729. 10.1101/849729

Verburgh, L., Scherder, E. J. A., van Lange, P. a. M., & Oosterlaan, J. (2016). The key to success in elite athletes? Explicit and implicit motor learning in youth elite and non-elite soccer players. Journal of Sports Sciences, 34(18), 1782–1790. 10.1080/02640414.2015.1137344

Verplanken, B., & Roy, D. (2016). Empowering interventions to promote sustainable lifestyles: Testing the habit discontinuity hypothesis in a field experiment. Journal of Environmental Psychology, 45, 127–134. 10.1016/j.jenvp.2015.11.008

Verplanken, B., Walker, I., Davis, A., & Jurasek, M. (2008). Context change and travel mode choice: Combining the habit discontinuity and self-activation hypotheses. Journal of Environmental Psychology, 28(2), 121–127. 10.1016/j.jenvp.2007.10.005

Virag, M., Janacsek, K., Horvath, A., Bujdoso, Z., Fabo, D., & Nemeth, D. (2015). Competition between frontal lobe functions and implicit sequence learning: Evidence from the long-term effects of alcohol. Experimental Brain Research, 233(7), 2081–2089. 10.1007/s00221-015-4279-8

Wagenmakers, E.-J., Wetzels, R., Borsboom, D., & van der Maas, H. L. J. (2011). Why psychologists must change the way they analyze their data: The case of psi: Comment on Bem (2011). Journal of Personality and Social Psychology, 100(3), 426–432. 10.1037/a0022790

Walker, I., Thomas, G. O., & Verplanken, B. (2015). Old Habits Die Hard: Travel Habit Formation and Decay During an Office Relocation. Environment and Behavior, 47(10), 1089–1106. 10.1177/0013916514549619

Wood, W., & Rünger, D. (2016). Psychology of Habit. Annual Review of Psychology, 67(Volume 67, 2016), 289–314. 10.1146/annurev-psych-122414-033417

Wood, W., Tam, L., & Witt, M. G. (2005). Changing circumstances, disrupting habits. Journal of Personality and Social Psychology, 88(6), 918–933. 10.1037/0022-3514.88.6.918

Woods, S. P., Scott, J. C., Sires, D. A., Grant, I., Heaton, R. K., Tröster, A. I., & HIV Neurobehavioral Research Center Group. (2005). Action (verb) fluency: Test-retest reliability, normative standards, and construct validity. Journal of the International Neuropsychological Society: JINS, 11(4), 408–415.

