## Supplementary material for "The interplay between executive functions and updating predictive representations"

| | Sham (n = 27, 15 females) | | Control (n = 32, 17 females) | | |
|---|---|---|---|---|---|
| | ***Mean*** | ***SD*** | ***Mean*** | ***SD*** | ***p-value (T-test)*** |
| **Age (years)** | 22.74 | 3.19 | 22.31 | 2.71 | 0.59 |
| **Education (years)** | 15.44 | 1.89 | 15.25 | 1.41 | 0.66 |
| **Learning** | 5.46 | 7.15 | 2.74 | 9.75 | 0.22 |
| **Rewiring** | 3.45 | 6.89 | 1.29 | 7.76 | 0.26 |
| **Retrieval Seq.A** | 4.67 | 7.59 | 2.03 | 9.17 | 0.23 |
| **Retrieval Seq. B** | 4.67 | 7.39 | 5.07 | 9.73 | 0.86 |
| **ANT alerting** | 33.89 | 31.99 | 38.73 | 53.10 | 0.67 |
| **ANT orienting** | 72.53 | 58.51 | 76.93 | 40.57 | 0.74 |
| **ANT executive** | 115.49 | 67.67 | 117.48 | 80.28 | 0.92 |
| **BCST** | 89.29 | 3.38 | 88.43 | 3.76 | 0.36 |
| **Counting Span** | 3.96 | 1.04 | 4.07 | 0.99 | 0.68 |
| **Go NoGo** | 3.31 | 0.40 | 3.36 | 0.46 | 0.70 |
| **Lexical fluency** | 15.26 | 4.18 | 14.63 | 3.92 | 0.55 |
| **Semantic fluency** | 23.00 | 4.87 | 25.47 | 5.41 | 0.07 |
| **Action fluency** | 21.11 | 7.13 | 20.03 | 5.56 | 0.53 |

**Supplementary Table 1.** Descriptive statistics and T-test comparisons for the Control and Sham groups. Participants in the current study took part in an experiment with multiple experimental groups, some of which received brain stimulation during the second session. We used data only from participants that did not receive verum stimulation (no stimulation -control- and sham). No differences between groups emerged in any of our measures.

| Predictor | B | β | SE B | t | p | 95% CI (Lower) | 95% CI (Upper) |
|---|---|---|---|---|---|---|---|
| (Intercept) | -18.106 | | 26.584 | -0.681 | 0.499 | -71.475 | 35.263 |
| ANT_executive | 0.019 | 0.187 | 0.013 | 1.422 | 0.161 | -0.008 | 0.045 |
| BCST_pers | 0.040 | 0.019 | 0.264 | 0.151 | 0.880 | -0.490 | 0.570 |
| CSPAN | 1.194 | 0.162 | 0.955 | 1.250 | 0.217 | -0.724 | 3.111 |
| GoNoGo | 5.881 | 0.341 | 2.203 | 2.669 | **0.010** | 1.458 | 10.304 |
| Lex_fluency | 0.154 | 0.084 | 0.256 | 0.602 | 0.550 | -0.360 | 0.669 |
| Sem_fluency | -0.455 | -0.325 | 0.191 | -2.390 | **0.021** | -0.838 | -0.073 |
| Act_fluency | -0.048 | -0.041 | 0.182 | -0.261 | 0.795 | -0.413 | 0.318 |

$R^2$ = 0.319, Adjusted $R^2$ = 0.226, F( 7 , 51 ) = 3.418 , p = 0.005

**Supplementary Table 2.** Multiple Regression Analysis of rewiring in Session 2 on executive function (EF) scores. The combination of EF scores explained 22.6% of the variance in rewiring GoNoGo and semantic fluency scores were the main predictors of our model with GoNoGo showing a positive relationship and semantic fluency showing a negative relationship with rewiring.

| Predictor | B | β | SE B | t | p | 95% CI (Lower) | 95% CI (Upper) |
|---|---|---|---|---|---|---|---|
| (Intercept) | -40.243 | | 34.392 | -1.170 | 0.247 | -109.288 | 28.802 |
| ANT_executive | 0.020 | 0.173 | 0.017 | 1.196 | 0.237 | -0.014 | 0.054 |
| BCST_pers | 0.371 | 0.153 | 0.342 | 1.087 | 0.282 | -0.315 | 1.057 |
| CSPAN | 0.987 | 0.114 | 1.236 | 0.798 | 0.428 | -1.494 | 3.468 |
| GoNoGo | 4.489 | 0.221 | 2.850 | 1.575 | 0.121 | -1.233 | 10.211 |
| Lex_fluency | 0.058 | 0.027 | 0.332 | 0.174 | 0.863 | -0.608 | 0.724 |
| Sem_fluency | -0.230 | -0.139 | 0.246 | -0.932 | 0.356 | -0.725 | 0.265 |
| Act_fluency | -0.259 | -0.188 | 0.236 | -1.101 | 0.276 | -0.733 | 0.214 |

$R^2$ = 0.177, Adjusted $R^2$ = 0.064, F( 7 , 51 ) = 1.57 , p = 0.166

**Supplementary Table 3.** Multiple Regression Analysis of learning in Session 1 on EF scores. The analysis yielded a non-significant model suggesting that the selected EF measures did not predict implicit statistical learning during the Learning Phase.

| Predictor | B | β | SE B | t | p | 95% CI (Lower) | 95% CI (Upper) |
|---|---|---|---|---|---|---|---|
| (Intercept) | -19.482 | | 35.371 | -0.551 | 0.584 | -90.491 | 51.528 |
| ANT_executive | 0.010 | 0.090 | 0.017 | 0.592 | 0.556 | -0.025 | 0.045 |
| BCST_pers | 0.201 | 0.085 | 0.351 | 0.572 | 0.570 | -0.504 | 0.906 |
| CSPAN | 0.158 | 0.019 | 1.271 | 0.125 | 0.901 | -2.393 | 2.710 |
| GoNoGo | 3.141 | 0.158 | 2.931 | 1.072 | 0.289 | -2.744 | 9.026 |
| Lex_fluency | 0.086 | 0.041 | 0.341 | 0.252 | 0.802 | -0.599 | 0.771 |
| Sem_fluency | -0.289 | -0.179 | 0.253 | -1.141 | 0.259 | -0.798 | 0.220 |
| Act_fluency | -0.083 | -0.061 | 0.242 | -0.343 | 0.733 | -0.570 | 0.403 |

$R^2$ = 0.092, Adjusted $R^2$ = -0.033, F( 7 , 51 ) = 0.736 , p = 0.642

**Supplementary Table 4.** Multiple Regression Analysis of retrieval of Sequence A in Session 3 on EF scores. The analysis yielded a non-significant model suggesting that the selected EF measures did not predict the retrieval of the initial knowledge in the Retrieval Phase.

| Predictor | B | β | SE B | t | p | 95% CI (Lower) | 95% CI (Upper) |
|---|---|---|---|---|---|---|---|
| (Intercept) | -50.028 | | 34.810 | -1.437 | 0.157 | -119.912 | 19.856 |
| ANT_executive | 0.016 | 0.141 | 0.017 | 0.961 | 0.341 | -0.018 | 0.051 |
| BCST_pers | 0.410 | 0.169 | 0.346 | 1.184 | 0.242 | -0.285 | 1.104 |
| CSPAN | 0.496 | 0.057 | 1.251 | 0.396 | 0.693 | -2.015 | 3.007 |
| GoNoGo | 5.739 | 0.284 | 2.885 | 1.989 | 0.052 | -0.052 | 11.531 |
| Lex_fluency | 0.065 | 0.030 | 0.336 | 0.193 | 0.847 | -0.609 | 0.739 |
| Sem_fluency | -0.040 | -0.024 | 0.249 | -0.161 | 0.873 | -0.541 | 0.461 |
| Act_fluency | -0.220 | -0.160 | 0.239 | -0.923 | 0.360 | -0.699 | 0.259 |

$R^2$ = 0.151, Adjusted $R^2$ = 0.034, F( 7 , 51 ) = 1.296 , p = 0.271

**Supplementary Table 5.** Multiple Regression Analysis of retrieval of Sequence B in Session 3 on EF scores. The analysis yielded a non-significant model suggesting that the selected EF measures did not predict the retrieval of the rewired knowledge in the Retrieval Phase.
